# Acid Ceramidase Inhibitor LCL-805 Antagonizes Akt Signaling and Promotes Iron-Dependent Cell Death in Acute Myeloid Leukemia

**DOI:** 10.1101/2023.10.21.563437

**Authors:** Johnson Ung, Su-Fern Tan, Todd E Fox, Jeremy JP Shaw, Maansi Taori, Bethany J Horton, Upendarrao Golla, Arati Sharma, Zdzislaw M Szulc, Hong-Gang Wang, Charles E. Chalfant, Myles C Cabot, David F Claxton, Thomas P Loughran, David J Feith

**Affiliations:** Department of Microbiology/Immunology/Cancer Biology, University of Virginia School of Medicine, Charlottesville, VA, USA; Department of Medicine, Division of Hematology & Oncology, University of Virginia School of Medicine, Charlottesville, VA, USA; University of Virginia Cancer Center, University of Virginia School of Medicine, Charlottesville, VA, USA; Department of Pharmacology, University of Virginia School of Medicine, Charlottesville, VA, USA; Department of Public Health Sciences, Division of Translational Research and Applied Statistics, University of Virginia School of Medicine, Charlottesville, VA, USA; Department of Medicine, Division of Hematology and Oncology, Pennsylvania State University College of Medicine, Hershey, PA, USA; Department of Pharmacology, Pennsylvania State University College of Medicine, Hershey, PA, USA; Penn State Cancer Institute, Pennsylvania State University College of Medicine, Hershey, PA, USA; Department of Biochemistry and Molecular Biology, Medical University of South Carolina College of Medicine, Charleston, SC, USA; Research Service, Richmond Veterans Administration Medical Center, Richmond VA, USA; East Carolina Diabetes and Obesity Institute, East Carolina University, Greenville, NC, USA; Department of Biochemistry & Molecular Biology, Brody School of Medicine, East Carolina University, Greenville, NC, USA

**Author notes:** Co-correspondence (Johnson Ung, David J Feith).

## Abstract

Acute myeloid leukemia (AML) is an aggressive hematologic malignancy requiring urgent treatment advancements. Ceramide is a cell death-promoting signaling lipid that plays a central role in therapy-induced cell death. Acid ceramidase (AC), a ceramide-depleting enzyme, is overexpressed in AML and promotes leukemic survival and drug resistance. The ceramidase inhibitor B-13 and next-generation lysosomal-localizing derivatives termed dimethylglycine (DMG)-B-13 prodrugs have been developed but remain untested in AML. Here, we report the *in vitro* anti-leukemic efficacy and mechanism of DMG-B-13 prodrug, LCL-805, across AML cell lines and primary patient samples. LCL-805 inhibited AC enzymatic activity, increased total ceramides, and reduced sphingosine levels. A median EC50 value of 11.7 μM was achieved for LCL-805 in cell viability assays across 32 human AML cell lines. As a single agent tested across a panel of 71 primary AML patient samples, a median EC50 value of 15.8 μM was achieved. Exogenous ceramide supplementation with C6-ceramide nanoliposomes, which is entering phase I/II clinical trial for relapsed/refractory AML, significantly enhanced LCL-805 killing. Mechanistically, LCL-805 antagonized Akt signaling and led to iron-dependent cell death distinct from canonical ferroptosis. These findings elucidated key factors involved in LCL-805 cytotoxicity and demonstrated the potency of combining AC inhibition with exogenous ceramide.

## 1. Introduction

Acute myeloid leukemia (AML) is an aggressive, heterogenous malignancy that afflicts tens of thousands of people annually [1]. Characterized by the clonal expansion of immature myeloid blasts, clinical features of AML include pancytopenia, anemia, and bone marrow failure [2]. Although individuals of all ages develop AML, the median age at diagnosis is 65 [3]. Intensive induction chemotherapy has been the primary treatment option since the 1970s with several targeted therapeutics approved in the 2010s [4]. Despite these approvals to supplement chemotherapy, poor treatment responses and high relapse rates persist. Moreover, treatment options are limited for elderly individuals that are unable to tolerate intensive induction chemotherapy.

Ceramides reside at the putative center of the complex sphingolipid metabolic pathway. Ceramide levels increase in response to chemotherapy [5], radiation [6], or targeted therapeutics [7] and contribute to treatment-induced cytotoxicity [8]. Multiple enzymes catabolize ceramide or modify its structure to generate increasingly complex sphingolipids. These metabolic paths reduce the ceramide pool to promote leukemia survival and drug resistance [9]. The development of ceramide-generating therapeutics is an enticing antineoplastic strategy as ceramides are important mediators of cell death and cancer cells modulate ceramide catabolism to maintain survival. Elevating intracellular ceramides through exogenous supplementation [10, 11] or metabolic blockade [12, 13] are actively being explored as anticancer therapies.

Aberrant sphingolipid metabolism is well documented in AML [14]. Our group previously identified and validated acid ceramidase (AC) as a therapeutic target [12]. AC is a lysosomal lipid hydrolase that catabolizes ceramides into sphingosine and free fatty acids. AC overexpression in AML is associated with and promotes chemotherapy resistance [15, 16]. AC targeting has been extensively studied in solid cancers with several inhibitors in preclinical development [17, 18]. B-13 is a ceramidase inhibitor with demonstrated cytotoxicity in multiple solid cancers [19, 20]. Unfortunately, the neutrophilic properties of B-13 preclude its accumulation in the lysosome where AC resides. To overcome this hurdle, next generation lysosome-localizing B-13 analogs termed dimethylglycine (DMG)-B-13 prodrugs were developed [21].

In the present study, we characterized the *in vitro* anti-leukemic efficacy and mechanisms of a DMG-B-13 prodrug, LCL-805, in AML for the first time. Our findings demonstrated that LCL-805 is an effective AC inhibitor and is cytotoxic toward AML cell lines and patient samples. LCL-805 cytotoxicity was further amplified via C6-ceramide nanoliposome (CNL) supplementation. Mechanistically, LCL-805 inhibited Akt signaling and induced iron-dependent cell death distinct from canonical ferroptosis.

## 2. Materials and Methods

### 2.1. Reagents

Inhibitors used for rescue studies: zVAD-FMK (Selleck Chemicals, #S7023); chloroquine (Selleck Chemicals, #S6999); bafilomycin A1 (Selleck Chemicals, #S1413); SC-79 (Cayman, #14972); deferoxamine mesylate (Selleck Chemicals, #S5742); liproxstatin-1 (Selleck Chemicals, #S7699); ferrostatin-1 (Selleck Chemicals, #S7243); GSK’872 (Selleck Chemicals, #S8465); necrostatin-1 (Selleck Chemicals, #S8037); pepstatin A (Cayman, #9000469); CA-074me (Selleck Chemicals, #S7420).

Anti-acid ceramidase antibody (#612302) was obtained from BD Transduction Laboratories. The following antibodies were obtained from Cell Signaling Technologies: caspase-3 (#9662); PARP (#9542); Mcl-1 (#5453); Bcl-2 (#4223); Bcl-xL (#2764); XIAP (#2042); LC3A (#4599); phospho-Akt (Ser473; #4060); Pan Akt (#4691); GPX4 (#52455); HIF-1α (#14179) β-actin (#3700); anti-rabbit IgG HRP-linked (#7074); anti-mouse IgG, HRP-linked (#7076).

### 2.2. Cell culture

Cell lines were cultured in a 37°C humidified incubator at 5% CO_2_. All cell lines were utilized within 6-8 weeks of thawing. OCI-AML3 were cultured in RPMI-1640 (Corning, #10-040) supplemented with 15% FBS (VWR, #97068–085). MOLM-16, F-36p, SKM-1, ME-1, SET-2, Kasumi-1, Kasumi-3, and Kasumi-6 were cultured in RPMI-1640 supplemented with 20% FBS. KG-1, KG-1/ABT737R, KG-1a, and KG-1a/ABT737R were cultured in IMDM (ThermoFisher #12440) supplemented with 20% FBS. MV4-11 were cultured in IMDM supplemented with 10% FBS. OCI-AML4 was cultured in α-MEM (ThermoFisher #12571063) supplemented with 20% FBS. Some cell lines required additional hGM-CSF supplementation (Miltenyi Biotec, #130–095-372): OCI-AML4 (100 ng/mL); F-36p, SKNO-1, and Kasumi-6 (10 ng/mL); and TF-1 (2 ng/mL). All remaining cell lines were cultured in RPMI-1640 supplemented with 10% FBS.

Drug-resistant cell lines were maintained in the presence of their respective drugs during routine cell culture. HL-60/VCR [22] were supplemented with 1 μg/ml vincristine sulfate (Cayman, #11764). ABT737 (Cayman, #11501) was supplemented to ABT737R cell lines: HL-60/ABT737R [23] (5 μM ABT737); KG-1/ABT737R and KG-1a/ABT737R [23] (1 μM ABT737).

Cell lines obtained from DSMZ include SKNO-1, OCI-M2, and SKM-1. All remaining cell lines were gifted (see acknowledgments) or acquired from ATCC. Short tandem repeat DNA profiling (LabCorp, formerly known as Genetica Cell Line Testing) was utilized to authenticate cell lines. All cell lines were confirmed negative for mycoplasma contamination using the MycoAlert PLUS kit (Lonza, #LT07–710). Non-targeting control (NT CT), ATG5 knockout (ATG5KO), and ATG7 knockout (ATG7KO) gene-edited MV4-11 cells were generated as previously described [24].

### 2.3 Primary AML patient samples and health normal samples

AML patient samples were collected with informed consents signed for sample collection according to approved protocols by the institutional review boards of the Milton S. Penn State Hershey Medical Center and the University of Virginia School of Medicine and in accordance with the Declaration of Helsinki (Supplemental Table 1). Normal PBMCs and bone marrow cells were purchased from AllCells (Alameda, CA), Virginia Blood Services, Zen-Bio (Durham, NC), and Inova Blood Donor Services (Sterling, VA). PBMCs and BM mononuclear cells were isolated from patients’ whole blood using the Ficoll-Paque gradient separation method and cryopreserved. Cryopreserved normal and AML patient PBMC samples were thawed and resuspended in RPMI growth medium with 15% FBS. Cell viability was determined using Muse Count & Viability Kit (Cytek MCH100102).

### 2.4. LCL-805 synthesis

LCL-805, dimaleate salt of (1R,2R)-2-N-(tetradecanoylamino)-1-(4’-nitrophenyl)-propyl-1,3-O-di-(N,N-dimethylamino) acetate, was prepared according to the previously published method from (1R,2R)-2-N-(tetradecanoylamino)-1-(4’-nitrophenyl)-propyl-1,3-diol and N,N-dimethylamino glycine, where in lieu of hydrochloric acid, maleic acid was applied in the final step [21].

### 2.5. Ceramide nanoliposome (CNL) formulation

Lipids, in chloroform, were combined at 3.75:1.75:0.75:0.75:3.0 molar ratio consisting of 1,2-dioleoyl-sn-glycero-3-phosphocholine/1,2-dioleoyl-sn-glycero-3-phosphoethanolamine/1,2-distearoyl-sn-glycero-3-phosphoethanolamine-N-[methoxy PEG(2000)]/PEG(750)-C8-ceramide/C6-ceramide. Lipids were dried under a stream of nitrogen gas. Lipids were subsequently hydrated with sterile 0.9% NaCl solution, sonicated, and underwent extrusion through 100 nm polycarbonate membranes at 60°C. CNL was sonicated in a 37°C water bath for 10-15 minutes before use.

### 2.6. Acid ceramidase (AC) activity assay

20,000 cells were seeded onto black 96-well assay plates (Corning, #3603). LCL-805 was prepared by diluting 1:10 in 37°C RPMI-1640 supplemented with 10% FBS. Cells were treated with vehicle (DMSO) or LCL-805 at the indicated concentrations in a final volume of 50 μL and incubated for the indicated times. After LCL-805 treatment, 50 μL of the AC substrate, RBM14C12 (Avanti Polar, #860855), was prepared in RPMI-1640 supplemented with 10% FBS and added to the cells at a final concentration of 16 μM and incubated for 3 hours. After incubation, 50 μL of 100% methanol was added followed immediately with 100 μL of 2.5 mg/ml sodium periodate in 100 mM glycine, pH 10.6. The plates were incubated for 2 hours at 37°C. After incubation, fluorescence was measured using the Synergy HT plate reader (Biotek, Winooski, VT; gain = auto; 365nm excitation/410-460nm emission). Following background subtraction, fluorescence was normalized to vehicle control, which was defined as 100%.

### 2.7. Immunoblotting

Cells were seeded between 5×10^5^ and 1×10^6^ cells/mL and treated with the indicated concentrations for the indicated timepoints. Following treatment, cells were pelleted, washed in 1X PBS, pelleted, resuspended in 1X RIPA buffer (Sigma, #R0278-500ML) supplemented with a protease inhibitor cocktail (Sigma, #P8340), phosphatase inhibitor cocktail 2 (Sigma, #P5726), and phosphatase inhibitor cocktail 3 (Sigma, #P0044), vortexed, and incubated on ice for 30 minutes. Samples were centrifuged at 16,000xg for 10 minutes at 4°C. Supernatant was collected and stored at -80°C. Protein levels were quantified using the bicinchoninic acid (BCA) protein assay kit (Pierce, #23225) following the manufacturer’s protocol. Samples were prepared in sample buffer (Invitrogen, #NP0007) and reducing agent (Invitrogen, #NP0009), denatured at 90°C for 10 minutes, centrifuged, and cooled at 4°C. Protein samples were resolved on a 4-12% Bis-Tris Plus 10-well gels (Invitrogen, #MW04120BOX). Proteins were transferred to LF-PVDF membranes (BioRad, #10026934) activated in 100% methanol. Following transfer, membranes were reactivated in 100% methanol.

Primary antibodies were diluted 1:1000 in 5% nonfat milk or 5% BSA dissolved in 1X TBS + 1% Tween-20 (TBST) as per the manufacturer protocol. Secondary antibodies were diluted 1:3333 in 5% nonfat milk dissolved in 1X TBST. Membranes were blocked for 1 hour at room temperature in 5% nonfat milk or 5% BSA dissolved in 1X TBST as per the manufacturer protocol followed by incubation with the primary antibodies on a rocker overnight at 4°C. Membranes were washed three times in 1X TBST and incubated with the appropriate secondary antibody for 1 hour at room temperature. For detection, the membranes were incubated in Clarity Western ECL Substrate (BioRad, #170-5061) or SuperSignal West Femto Maximum Sensitivity Substrate (BioRad, #34096) and visualized by chemiluminescence using the BioRad ChemiDoc MP imaging system. Protein quantification was performed using Image Lab 6.0.1 software.

### 2.8. Sphingolipid profiling

Lipids were extracted from cell homogenates equivalent to 400 mg of protein using a mix of isopropanol:water:ethyl acetate (3:1:6; v:v:v). Internal standards (10 pmol of d17 long-chain bases and C12 acylated sphingolipids) were added to samples at the onset of the extraction procedure. Extracts were separated on a Waters I-class Acquity UPLC chromatography system. Mobile phases were (A) 60:40 water:acetonitrile and (B) 90:10 isopropanol:methanol with both mobile phases containing 5 mM ammonium formate and 0.1% formic acid. A Waters C18 CSH 2.1 mm ID × 10 cm column maintained at 65°C was used for the separation of the sphingoid bases, 1-phosphates, and acylated sphingolipids. The eluate was analyzed with an inline Waters TQ-S mass spectrometer using multiple reaction monitoring.

### 2.9. Cell line viability assays

20,000 cells were seeded onto 96-well assay plates (Corning, #353072). LCL-805 was prepared by creating a 1:10 working stock in 37°C RPMI-1640 supplemented with 10% FBS. Cells were treated with vehicle (DMSO) or LCL-805 at the indicated concentrations in a final volume of 100 μL and incubated for the indicated times. Following incubation, 20 μL of [3-(4,5-dimethylthiazol-2-yl)-5-(3-carboxymethoxyphenyl)-2-(4-sulfophenyl)-2H-tetrazolium] (MTS) (Promega, #G3582) were added and incubated at 37°C for 3 hours. Absorbance of the MTS-derived formazan product was measured at 490 nm with the Synergy HT plate reader (Biotek, Winooski, VT). Following background subtraction, absorbances were normalized to vehicle control, which was defined as 100%. EC50 values were calculated with GraphPad Prism v9.5.1 using the function: “log(inhibitor) vs. normalized response -- variable slope.

### 2.10. Patient sample viability assays

5,000 cells/well were seeded onto 384-well plates (ThermoFisher, #165195) and treated with vehicle (DMSO), LCL-805, CNL, or combination for 48 hours at the indicated concentrations. Viability was determined using the CellTiter-Glo luminescence assay (Promega, #G755B). Luminescence was measured using the GloMax Discover (Promega) as per the manufacturer’s protocol. Background media luminescence was subtracted from all samples and luminescence data from drug-treated conditions were normalized to DMSO vehicle, which was set to 100%. EC50 values were determined as in section 2.9.

### 2.11. Inhibitor rescue studies

For inhibitor rescue studies, cells were pretreated with vehicle (DMSO) or the indicated compounds for 2 hours. Concentrations for rescue inhibitors were based on prior publications [24, 25]. We utilized rescue inhibitor concentrations at, above, and below the published concentrations and observed similar effects for each concentration. Following preincubation, cells were utilized for MTS viability studies, immunoblotting, or flow cytometry as described in section 2.9, section 2.7, and section 2.12, respectively.

### 2.12. Flow cytometry

2.5×10^6^ cells were seeded at a density of 2.5×10^5^ cells/mL. LCL-805 was prepared by creating a 1:10 working stock in 37°C RPMI-1640 supplemented with 10% FBS. Cells were treated with vehicle (DMSO) or LCL-805 at the indicated concentrations and incubated for the indicated times. Following treatment, cell death and mitochondrial membrane potential were evaluated using Muse Annexin V & Dead Cell Kit (Cytek, #MCH100105) and Muse MitoPotential Kit (Cytek, #MCH100110) per the manufacturer’s protocol. For each assay, the final cell dilutions were 1:4 and 500-2000 events were captured.

### 2.13. Synergy analysis

Two drug synergy was quantified using the online SynergyFinder 2.0 software [26]. The following parameters were used during analysis: Readout = Viability; Detect Outliers = Yes; Curve Fitting = LL4. Synergy scores represent the percentage differences between observed responses versus predicted responses calculated using the Bliss reference model. Scores above 10, between 10 and -10, and below -10 are considered synergistic, additive, and antagonistic, respectively.

### 2.14. Statistical analysis

Indicated statistical tests were performed using GraphPad Prism version 10.0.2. Results presented are representative data of at least two independent experiments. Each experiment contained at least three technical replicates. Error bars represent standard deviation.

## 3. Results

### 3.1. LCL-805 inhibited AC in a concentration- and time-dependent manner

LCL-805 is an amino acid prodrug of B-13 and is structurally related to LCL-521 [21]. This class of prodrug traffics to lysosomes via its amino acid moiety where they are metabolized into the active ceramidase inhibitor, B-13 [21]. We utilized a fluorescence-based assay to evaluate the effect of LCL-805 on AC activity [27]. Treatment of human AML cell lines MM-6 and OCI-AML2 with LCL-805 significantly reduced AC enzymatic activity in a concentration- and time-dependent manner (**Figure 1A, B**). Detection of the 13 kDa α-AC subunit is frequently used as a marker of active AC enzyme [28]. Consistent with the results of our activity assays, LCL-805 reduced levels of the 13 kDa AC subunit in MM-6 and OCI-AML2 cells in a time-dependent manner (**Figure 1C**). Both readouts showed transient AC inhibition within 2 hours post LCL-805 treatment. Together, these results demonstrated the AC inhibitory effects of LCL-805.

**Figure 1.**
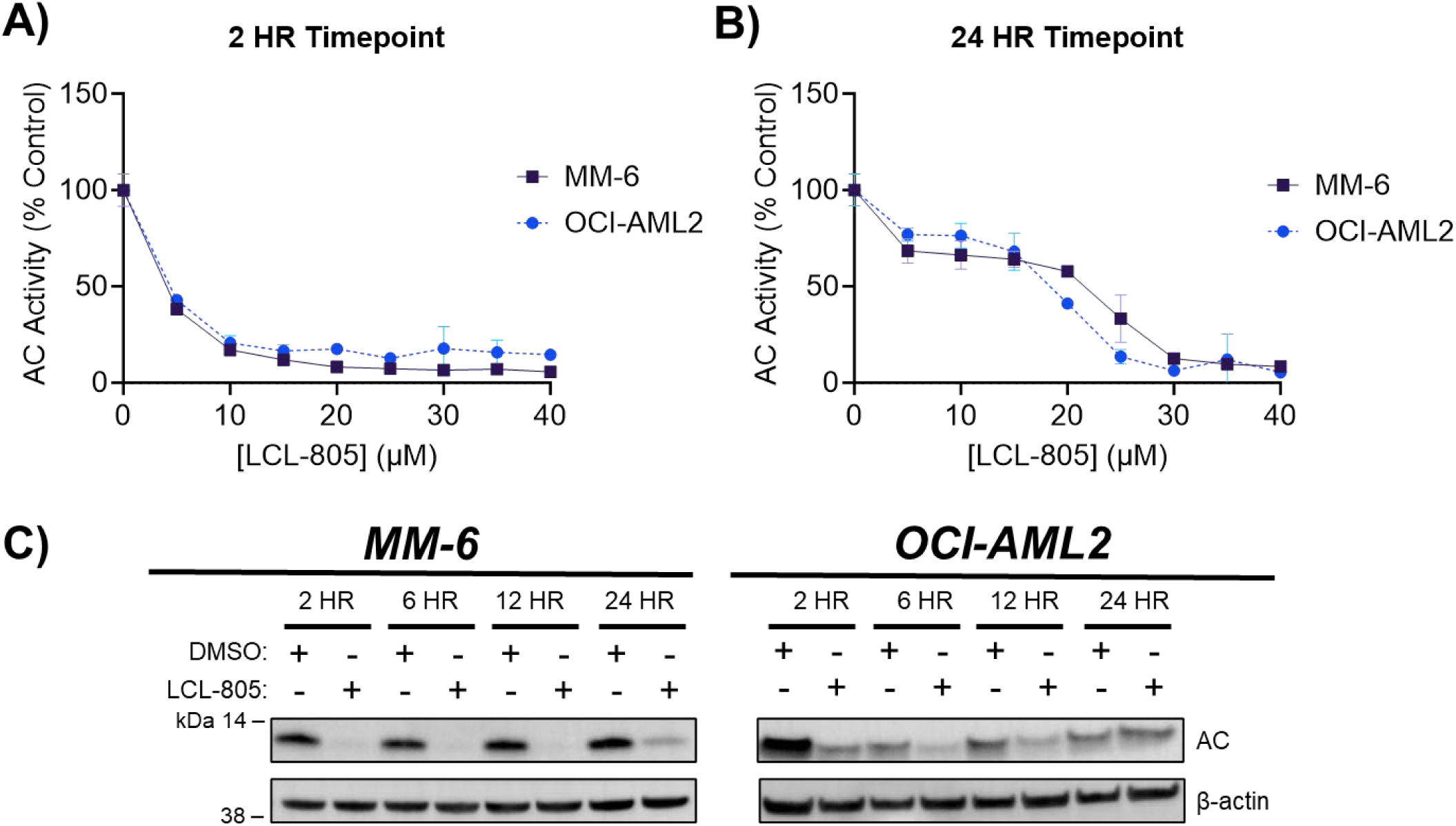
LCL-805 inhibited AC in a concentration- and time-dependent manner. (A,B) Fluorogenic AC activity assay conducted in MM-6 and OCI-AML2 human AML cell lines. Cells were treated with vehicle (DMSO) or the indicated concentration of LCL-805 for the indicated times. Error bars represent +/- standard deviation. (C) Immunoblotting of AC protein levels (α-subunit) in MM-6 and OCI-AML2 cells treated with DMSO or LCL-805 (15 µM) for the indicated times. Data presented are representative experiments with at least three technical replicates.

### 3.2. LCL-805 increased cellular ceramide and decreased sphingosine levels

We next measured the effect of LCL-805 on sphingolipid content at these same timepoints. AC is part of the sphingolipid metabolism “salvage pathway” and catabolizes ceramides into sphingosine and fatty acids. Upon AC inhibition, ceramides are expected to increase while sphingosine levels concurrently decrease. Indeed, sphingosine levels decreased substantially after 2 and 6 hours LCL-805 treatment in MM-6 and OCI-AML2 cells (**Figure 2A, B**). Sphingosine-1-phosphate (S1P) levels were minimally changed (**Figure S1**). Total ceramides were significantly elevated with the greatest increase observed in OCI-AML2 cells (**Figure 2C, D**). Long-chain ceramides, such as C16-ceramide, may be more cytotoxic than very long-chain ceramides, such as C24:1-ceramide [29]. We previously demonstrated that elevated C16/C24:1 ceramide ratios are associated with increased cell death [11]. LCL-805 increased both C16- and C24:1-ceramides (**Figure 2E, F**) and C16/C24:1 ceramide ratios (**Figure 2G, H**). Similar findings were observed for C16/C24 ceramide ratios (data not shown). Ceramide elevation and sphingosine depletion were rapid but diminished over time. Levels of all ceramide species are shown in Figure S2 as well as comprehensive profiling of additional ceramide metabolites (**Figures S3, S4, S5).** In summary, dihydroceramides were increased following LCL-805 treatment in both cell lines (**Figure S3**). Some hexosylceramides were increased (**Figure S4**) while sphingomyelin levels were mostly unchanged following LCL-805 treatment (**Figure S5**). Taken together, these results supported the AC-inhibitory effects of LCL-805 which resulted in increased ceramides at the expense of sphingosine.

**Figure 2.**
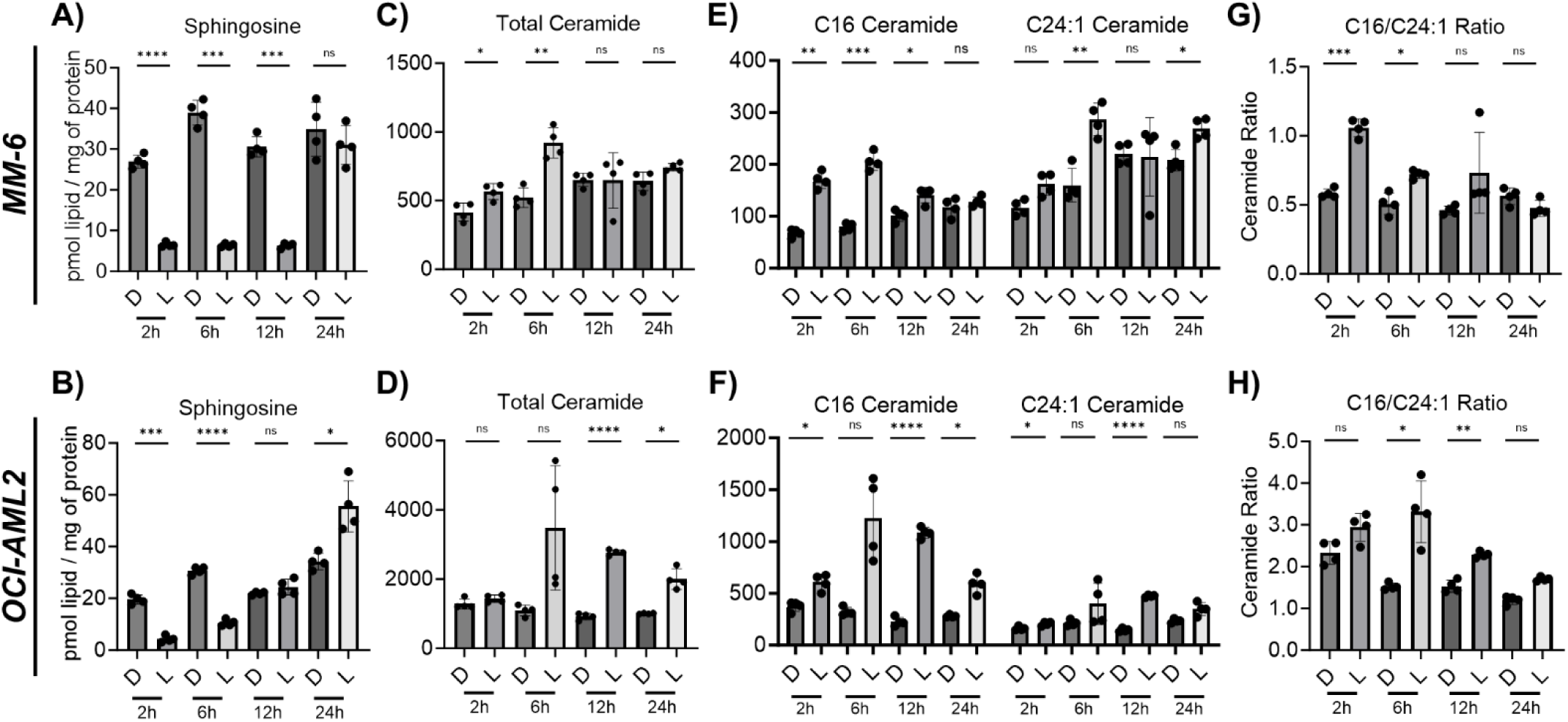
LCL-805 increased cellular ceramide and decreased sphingosine levels. Sphingolipid profiling of MM-6 and OCI-AML2 cells treated with vehicle (DMSO) or LCL-805 (15 µM) for the indicated times. Levels of sphingosine (A,B), total ceramides (C,D), C16 and C24:1 ceramides (E,F), and C16/C24:1 ceramide ratios (G,H) were evaluated by liquid chromatography mass spectrometry. Bars represent the average of four technical replicates from a representative experiment. Error bars represent +/- standard deviation. Statistical analyses represent Welch’s ANOVA with Dunnett’s T3 multiple comparisons test. * p < 0.05, ** p < 0.005, *** p < 0.001, **** p < 0.0001, ns = non-significant. D = DMSO; L = LCL-805. C## = fatty acid chain length. Total = sum of sphingolipid species.

### 3.3. LCL-805-induced loss of AML cell viability is characterized by increased phosphatidylserine externalization and mitochondrial depolarization

We next evaluated LCL-805 effects on cell viability across 32 human AML cell lines. A median EC50 of 11.7 μM was achieved across these cell lines (**Figure 3A, B**). We also utilized several parental/drug-resistant cell line pairs and showed that LCL-805 was equipotent in drug-resistant cell lines (**Figure 3B**). LCL-805 was then evaluated as a single agent across a panel of primary AML patient samples. LCL-805 demonstrated efficacy (median EC50 value of 15.8 μM) in 52.9% of the patient samples. The remaining 47.1% of the samples were resistant to the maximal concentration tested of 20 μM (**Figure 3C**).

**Figure 3.**
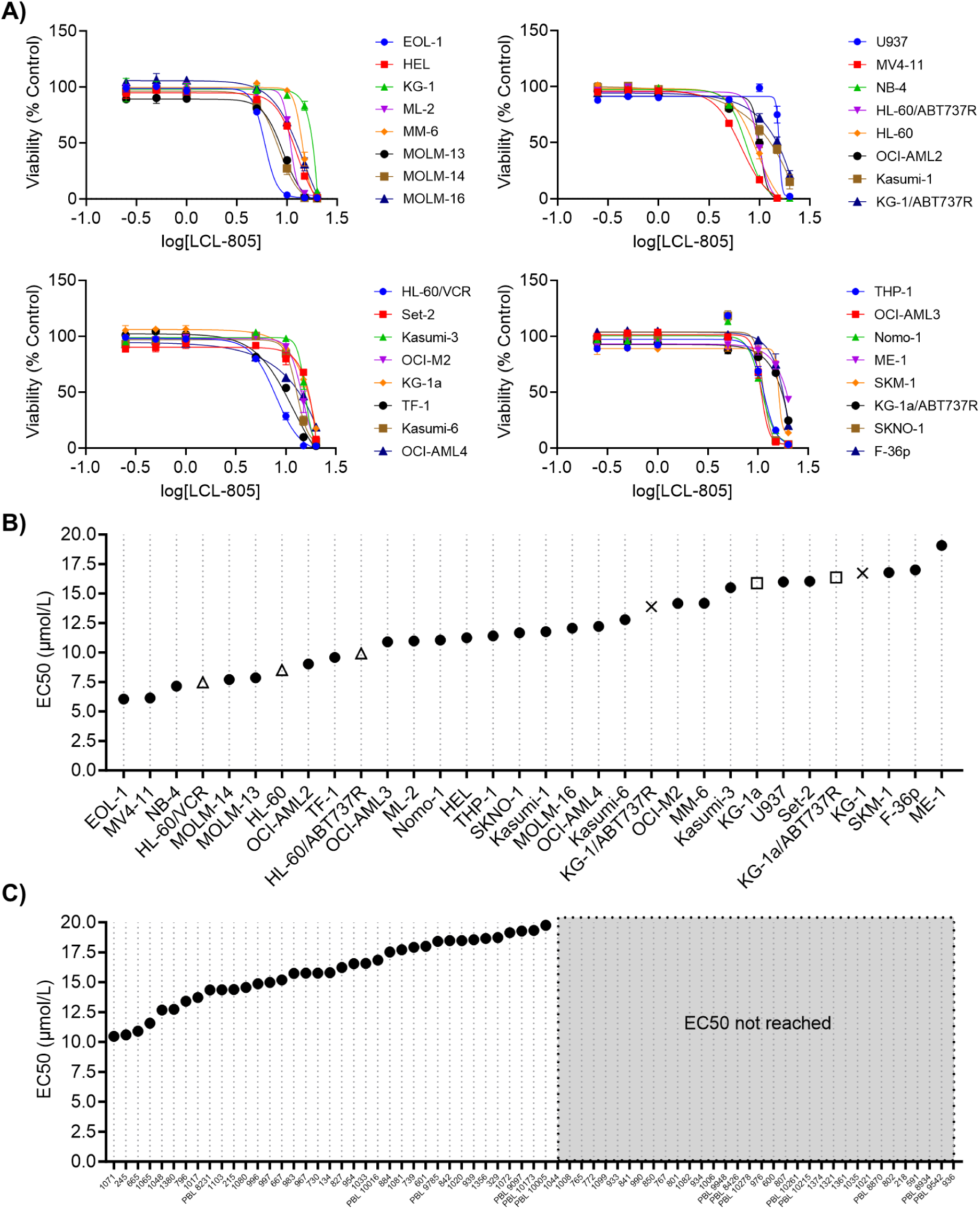
LCL-805 reduced viability of human AML cell lines and primary patient samples. (A) Cell viability of 32 human AML cell lines treated with vehicle (DMSO) or the indicated concentrations of LCL-805 for 48 hours. (B,C) Calculated LCL-805 EC50 values across AML cell lines (B) or primary AML patient samples (C). Legend for parental and drug-resistant cell line pairs: HL-60 (△); KG-1 (x); KG-1a (□); ABT737R = ABT737 resistant; VCR = vincristine resistant. Bars represent the average of three technical replicates from a representative experiment. Error bars represent +/- standard deviation.

We next utilized flow cytometry to measure phosphatidylserine externalization (annexin V staining) and 7-aminoactinomycin D (7-AAD staining). We observed a significant concentration-dependent increase in single annexin V positive, single 7-AAD-positive, and double positive cells following LCL-805 treatment (**Figure 4A, B)**. As AC inhibition is deleterious to mitochondrial function in AML [30] and pancreatic cancer [31], we investigated the effects of LCL-805 on mitochondrial membrane potential. In MM-6 and OCI-AML2 cells, LCL-805 treatment promoted concentration-dependent mitochondrial depolarization (**Figure 4C, D**). Taken together, these findings demonstrated that LCL-805 reduced cell viability, induced phosphatidylserine externalization, and triggered mitochondrial membrane depolarization within the micromolar range.

**Figure 4.**
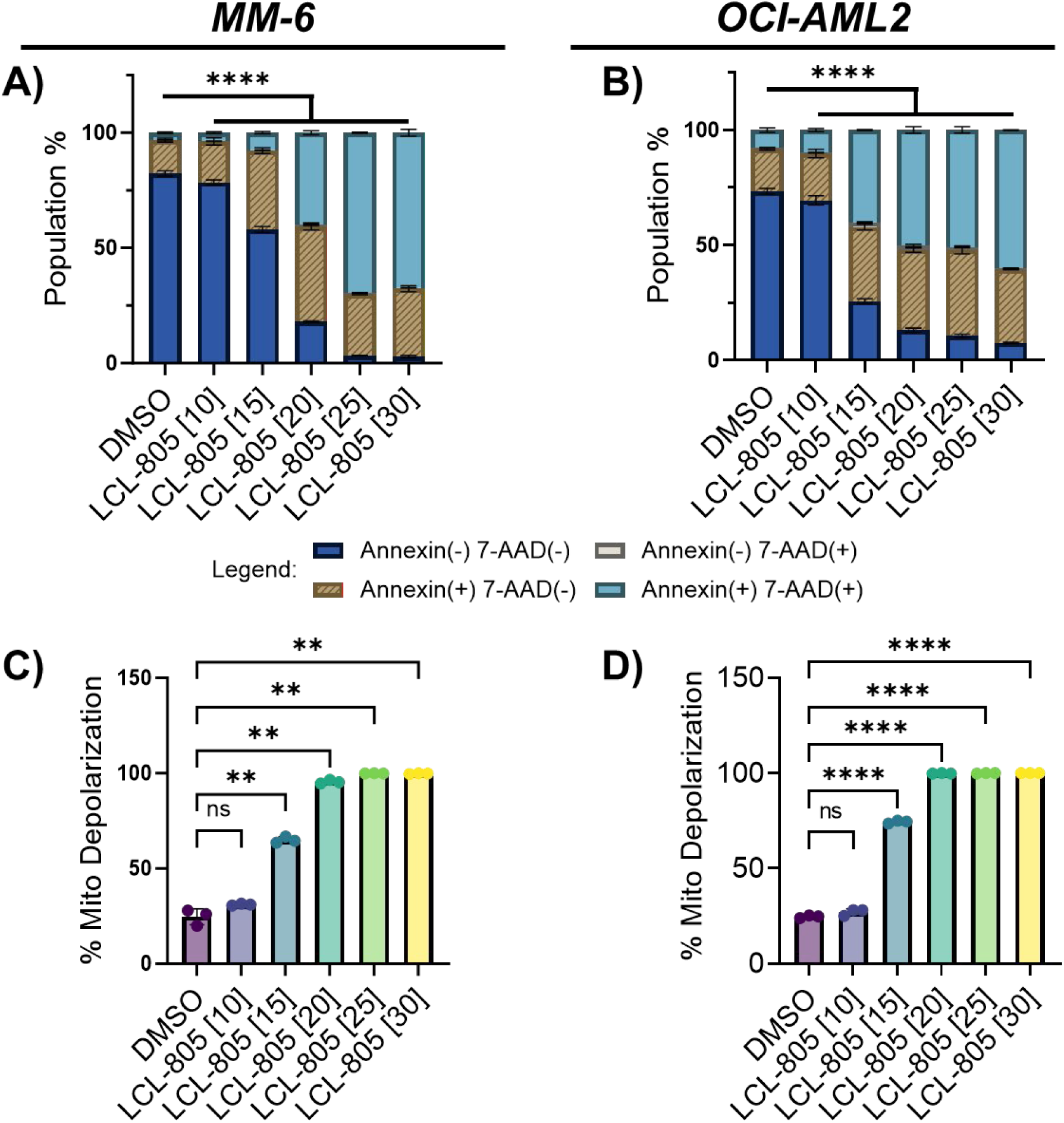
LCL-805 increased phosphatidylserine externalization and triggered mitochondrial depolarization. (A,B) Annexin V and 7-AAD flow cytometry in MM-6 and OCI-AML2 treated for 24 hours with vehicle (DMSO) or the indicated concentrations of LCL-805 (in µM, shown in brackets). (C,D) Mitochondrial membrane potential in MM-6 and OCI-AML2 treated for 24 hours with DMSO or the indicated concentrations of LCL-805. Bars represent the average of three technical replicates from a representative experiment. Error bars represent +/- standard deviation. Statistical analyses represent two-way ANOVA with Dunnett’s multiple comparisons test (A,B) and Welch’s ANOVA with Dunnett’s multiple comparisons test (C,D). ** *p* < 0.005, **** *p* < 0.0001, ns = non-significant.

### 3.4. Inhibition of caspases, lysosomal cathepsins, autophagy, or necroptosis did not protect against LCL-805 cytotoxicity

Previous reports of B-13 analog cell death mechanisms are divergent with some indicating caspase-dependent apoptosis [12] and others indicating caspase-independent non-apoptotic cell death [32]. To determine whether caspase-3 is activated following LCL-805 treatment, we treated AML cells with LCL-805 and measured caspase-3 cleavage via immunoblotting. We detected partial cleaved caspase-3 in MM-6 which increased over time. Weaker cleaved caspase-3 was detected in OCI-AML2 cells (**Figure 5A, B**). Poly (ADP-ribose) polymerase-1 (PARP) cleavage is routinely used as a marker of caspase activation and was observed at 12 hours and 24 hours post LCL-805 treatment, with weaker cleavage detected in OCI-AML2 cells (**Figure 5A, B**). Sphingolipids are linked to Bcl-2 family proteins, gatekeepers of caspase-dependent apoptotic cell death [14], and the endogenous caspase inhibitor, X-linked inhibitor of apoptosis protein (XIAP) [33, 34]. LCL-805 treatment did not decrease protein levels of the anti-apoptotic proteins Mcl-1, Bcl-2, Bcl-xL, or XIAP (**Figure 5A, B**). Pretreatment with the pan-caspase inhibitor, zVAD-FMK, did not protect MM-6 or OCI-AML2 cells from LCL-805 cytotoxicity (**Figure 6A, B**). As other iterations of B-13 analogs destabilize lysosomes [35] or induce cell death via lysosomal cathepsin activity [32], we also evaluated the contribution of lysosomal cathepsins in LCL-805 killing. Pretreating cells with CA-074me or pepstatin-A, cathepsin B and D inhibitors respectively, did not protect against LCL-805 killing (**Figure S6A, B**). These findings show that caspases or lysosomal cathepsins were not the primary mediators of LCL-805 cytotoxicity.

**Figure 5.**
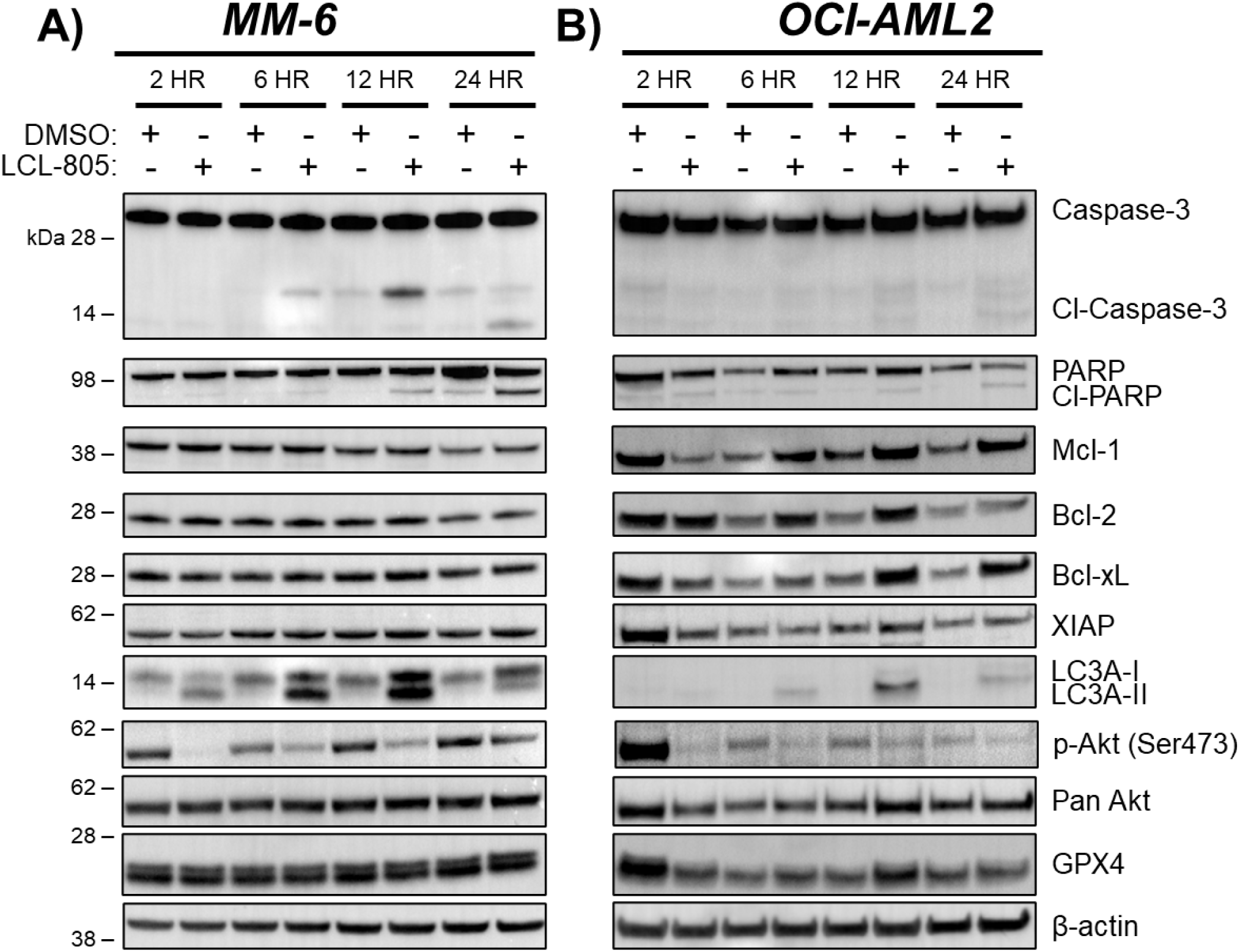
LCL-805 induced caspase-3 and PARP cleavage, LC3A processing, and reduced Akt phosphorylation. Immunoblotting of MM-6 (A) and OCI-AML2 cells (B) treated with DMSO or LCL-805 (15 µM) for the indicated times. Data are from a representative experiment.

**Figure 6.**
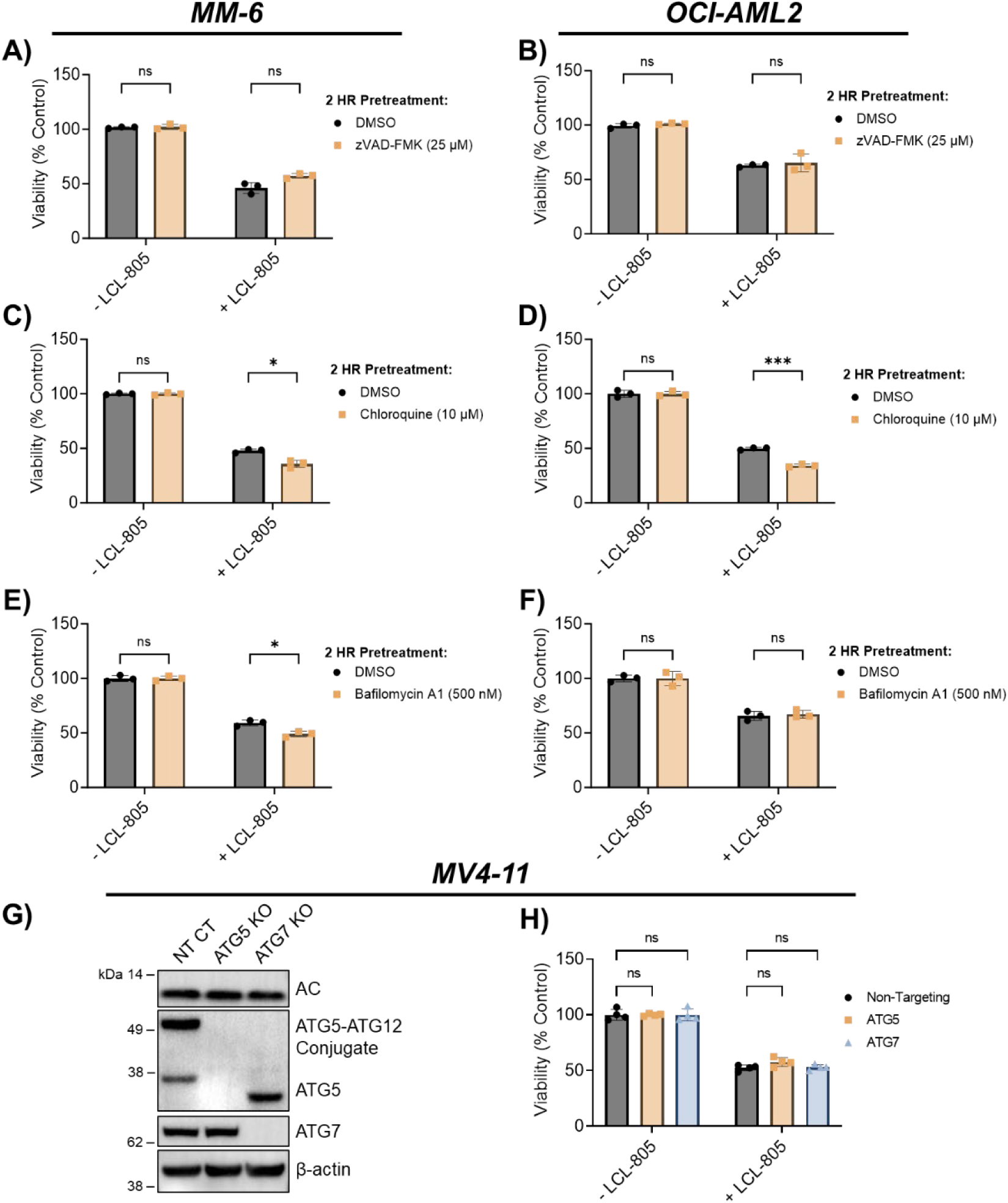
Inhibition of caspases or autophagy did not rescue LCL-805-mediated cell death. (A,B) Cell viability of MM-6 and OCI-AML2 cells pretreated with vehicle (DMSO) or zVAD-FMK (25 µM) for 2 hours and subsequently treated with DMSO or LCL-805 (15 µM) for 24 hours. (C,D) Cell viability of MM-6 and OCI-AML2 cells pretreated with vehicle (DMSO) or chloroquine (10 µM) for 2 hours and subsequently treated with DMSO or LCL-805 (C = 20 µM; D = 17.5 µM) for 24 hours. (E,F) Cell viability of MM-6 and OCI-AML2 cells pretreated with vehicle (DMSO) or bafilomycin A1 (500 nM) for 2 hours and subsequently treated with DMSO or LCL-805 (E = 20 µM; F = 17.5 µM) for 24 hours. (G) Immunoblotting of AC (α-subunit), ATG5, and ATG7 protein levels in non-targeting control (NT CT), ATG5-deficient (ATG5 KO), or ATG7-deficient (ATG7 KO) MV4-11 cells. (H) Cell viability of NT CT, ATG5 KO, or ATG7 KO MV4-11 cells treated with vehicle (DMSO) or LCL-805 (10 µM) for 24 hours. Bars represent the average of three technical replicates from a representative experiment. Error bars represent +/- standard deviation. Statistical analyses represent unpaired Welch’s t-tests with Holm-Šídák’s multiple comparisons test (A-F) and two-way ANOVA with Dunnett’s multiple comparisons test (H). * p < 0.05, *** p < 0.001, ns = non-significant.

We next evaluated whether autophagy was involved in LCL-805-mediated cell death. Microtubule-associated protein light chain 3 (LC3) is a key component of autophagosome formation and its processing from LC3-I to LC3-II is commonly used to measure autophagosome formation and autophagic flux [36]. Increased LC3A-II protein levels after LCL-805 treatment suggested increased autophagosome formation or decreased autophagic flux (**Figure 5A, B**). We leveraged pharmacological and genetic autophagy inhibition to evaluate the role of autophagy in LCL-805-mediated killing. Chloroquine and bafilomycin A1 are late-stage autophagy inhibitors that antagonize autophagosome fusion with the lysosome [37] and disrupt acidification of the autolysosome [38]. Pretreatment with chloroquine (**Figure 6C, D**) or bafilomycin A1 (**Figure 6E, F**) did not protect against but rather slightly enhanced LCL-805 toxicity. Autophagy-Related 5 (ATG5) and Autophagy-Related 7 (ATG7) are critical for autophagosome formation and LC3 processing [36]. To determine how depleting autophagy machinery affects LCL-805 killing, we evaluated LCL-805 toxicity in MV4-11 human AML cells deficient in ATG5 or ATG7 previously generated by our group [24]. Neither ATG5 nor ATG7 knockout affected AC protein levels (**Figure 6G**). Consistent with our pharmacological inhibition studies, knockout of ATG5 or ATG7 did not blunt LCL-805 cytotoxicity (**Figure 6H**). We next evaluated whether necroptotic cell death was involved in LCL-805 killing. Ceramides have previously been implicated in necroptosis, a form of regulated cell death that involves the activation of receptor-interacting protein kinase 1 (RIPK1) and receptor-interacting protein kinase 3 (RIPK3) [39, 40]. Pretreating AML cells with the RIPK1 inhibitor, necrostatin-1, or the RIPK3 inhibitor, GSK’872, did not protect against LCL-805 cytotoxicity (**Figure S6C**). Taken together, these findings suggested that autophagy and RIPK1/3 activation were not the primary mediators of LCL-805 cytotoxicity.

### 3.5. Akt reactivation and iron chelation rescued LCL-805-induced cell death

Several studies have associated AC with oncogenic Akt signaling [41, 42], which prompted us to evaluate the effects of LCL-805 on Akt. Akt phosphorylation at Ser473 and Thr308 contributes to its full activity. LCL-805 treatment significantly reduced Ser473 phosphorylation in MM-6 and OCI-AML2 cells (**Figure 5A, B**). We did not detect significant Akt phosphorylation at Thr308 in either cell line (data not shown). To assess whether Akt antagonism mediated LCL-805 cytotoxicity, we measured cell viability in cells pretreated with the Akt activator SC-79, which binds Akt and promotes an active conformation [43], then treated with LCL-805. SC-79 pretreatment protected against LCL-805-induced loss of cell viability in both MM-6 and OCI-AML2 cells (**Figure 7A, B**). In MM-6 cells, SC-79 pretreatment increased Ser473 Akt phosphorylation following 24-hour LCL-805 treatment. In OCI-AML2 cells, we observed weaker increases in Ser473 phosphorylation (**Figure 7C**). Together, these findings demonstrated that Akt inhibition contributed to LCL-805 toxicity and pharmacological Akt reactivation protected against LCL-805 cytotoxicity.

**Figure 7.**
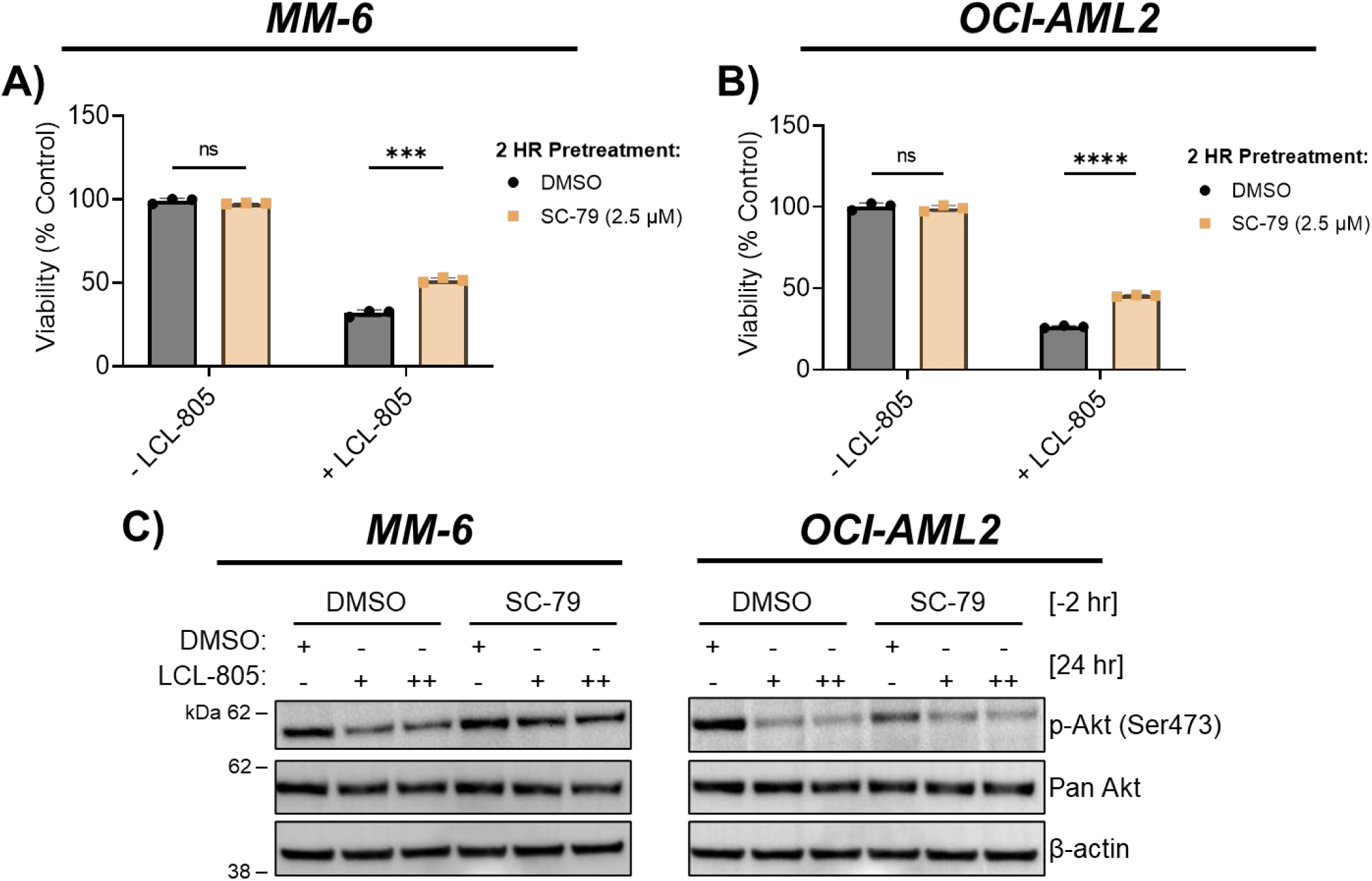
Akt reactivation rescued LCL-805-induced cell death. (A,B) Cell viability of MM-6 and OCI-AML2 cells pretreated with vehicle (DMSO) or SC-79 (2.5 µM) for 2 hours and subsequently treated with DMSO or LCL-805 (A = 20 µM; B = 17.5 µM) for 24 hours. (C) Immunoblotting of MM-6 and OCI-AML2 cells pretreated with SC-79 (2.5 µM) for 2 hours then treated with DMSO or LCL-805 (MM-6: + = 17.5 µM, ++ = 20 µM; OCI-AML2: + = 12.5 µM, ++ = 15 µM) for 24 hours. Bars represent the average of three technical replicates from a representative experiment. Error bars represent +/- standard deviation. Statistical analyses represent unpaired Welch’s t-tests and Holm-Šídák’s multiple comparisons test. *** *p* < 0.001, **** *p* < 0.0001, ns = non-significant.

As an extension of their role as cellular recycling centers, lysosomes are critical regulators of iron trafficking, storage, and metabolism [44]. Deferoxamine mesylate (DFO) is an FDA-approved iron chelating agent used to treat acute iron intoxication and chronic iron overload. We tested the effects of iron chelation on LCL-805 toxicity and found that DFO pretreatment significantly protected AML cells from LCL-805 cytotoxicity (**Figure 8 A, B**). Because iron is necessary for ferroptotic cell death, which is characterized by iron accumulation and lipid peroxidation, we evaluated whether pharmacological ferroptosis inhibitors rescued LCL-805 killing. Treatment with the ferroptosis inhibitors liproxstatin-1 (**Figure 8C, D**) or ferrostatin-1 (**Figure 8E, F**) did not attenuate LCL-805-mediated cell death. Moreover, LCL-805 treatment did not affect protein levels of glutathione peroxidase 4 (GPX4), a master regulator of canonical ferroptosis (**Figure 5A, B**). To determine how DFO protected against LCL-805 cytotoxicity, we evaluated protein changes in LCL-805-treated AML cells with or without DFO pretreatment (**Figure 8G**). GPX4 protein levels and Akt phosphorylation were unchanged following LCL-805 with or without DFO pretreatment. DFO is a known HIF-1α activator [45]. While treatment with DFO alone increased HIF-1α protein levels in MM-6 cells, DFO and LCL-805 treatment synergistically stabilized HIF-1α protein expression suggesting a protective role for HIF-1α in LCL-805 cytotoxicity. Taken together, these findings demonstrated that LCL-805 antagonized Akt signaling and induced iron-dependent cell death distinct from canonical ferroptosis.

**Figure 8.**
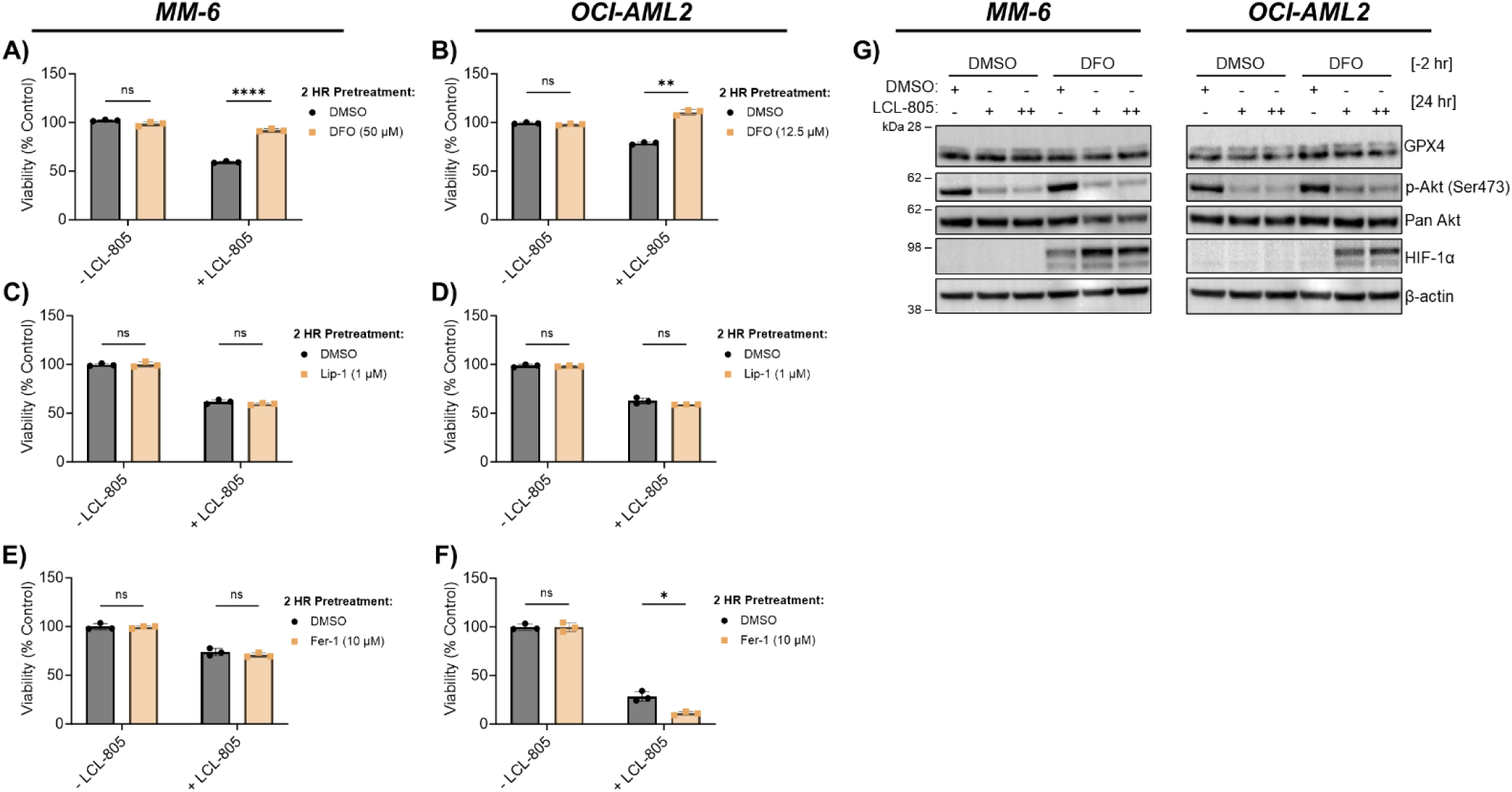
Iron chelation rescued LCL-805-induced cell death. (A,B) Cell viability of MM-6 and OCI-AML2 cells pretreated with vehicle (DMSO) or the indicated concentration of DFO for 2 hours and subsequently treated with DMSO or LCL-805 (15 µM) for 24 hours. (C,D) Cell viability of MM-6 and OCI-AML2 cells pretreated with vehicle (DMSO) or the indicated concentration of liproxstatin-1 (Lip-1) for 2 hours and subsequently treated with DMSO or LCL-805 (C = 20 µM; D = 17.5 µM) for 24 hours. (E,F) Cell viability of MM-6 and OCI-AML2 cells pretreated with vehicle (DMSO) or the indicated concentration of ferrostatin-1 (Fer-1) for 2 hours and subsequently treated with DMSO or LCL-805 (E = 15 µM; F = 15 µM) for 24 hours. (G) Immunoblotting of MM-6 and OCI-AML2 cells pretreated with DFO for 2 hours (DFO; MM-6 = 50 µM; OCI-AML2 = 12.5 µM) then treated with DMSO or LCL-805 (MM-6: + = 15 µM, ++ = 17.5 µM; OCI-AML2: + = 12.5 µM, ++ = 15 µM) for 24 hours. Bars represent the average of three technical replicates from a representative experiment. Error bars represent +/- standard deviation. Statistical analyses represent unpaired Welch’s t-tests and Holm-Šídák’s multiple comparisons test. * *p* < 0.05, ** *p* < 0.005, **** *p* < 0.0001, ns = non-significant.

### 3.6. C6-ceramide nanoliposome (CNL) supplementation improved LCL-805 toxicity

LCL-805 increased pro-death ceramides and the C16/C24:1 and C16/C24 ceramide ratios, which prompted us to evaluate the cytotoxic effects of combining exogenous ceramide with LCL-805. To this end, we tested LCL-805 and CNL across a panel of primary AML patient samples. The effect of LCL-805, CNL, or the combination on cell viability and the corresponding Bliss synergy scores from the top five patient samples most sensitive to the combination are shown in Figures 9A and 9B. As single agents, LCL-805 and CNL demonstrated modest efficacy. However, the combination resulted in highly potent synergistic lethality (**Figure 9A, B**). Treatment of AML patient sample with the CNL and LCL-805 combination demonstrated synergistic (n=48) or additive (n= 16) Bliss synergy scores while only one exhibited antagonism (**Figure 9C)**. We also compared these results in primary AML patient samples to nine healthy donor normal samples (seven PBMC and two whole bone marrow samples). The combination of LCL-805 and CNL reduced cell viability more in leukemic AML patient samples compared to healthy donor samples although the differences were not statistically significant (**Figure 9D**). Cell viability dropped to less than 50% in only 52.9% of patient samples treated with single agent LCL-805 up to 20 µM (**Figure 3C**). However, the addition of CNL (5 µM) with 15 µM LCL-805 resulted in loss of cell viability greater than 50% in 70.8% of patient samples (**Figure 9D**). Together, these results demonstrated the promising therapeutic effects of combining AC inhibition with exogenous ceramide supplementation.

**Figure 9.**
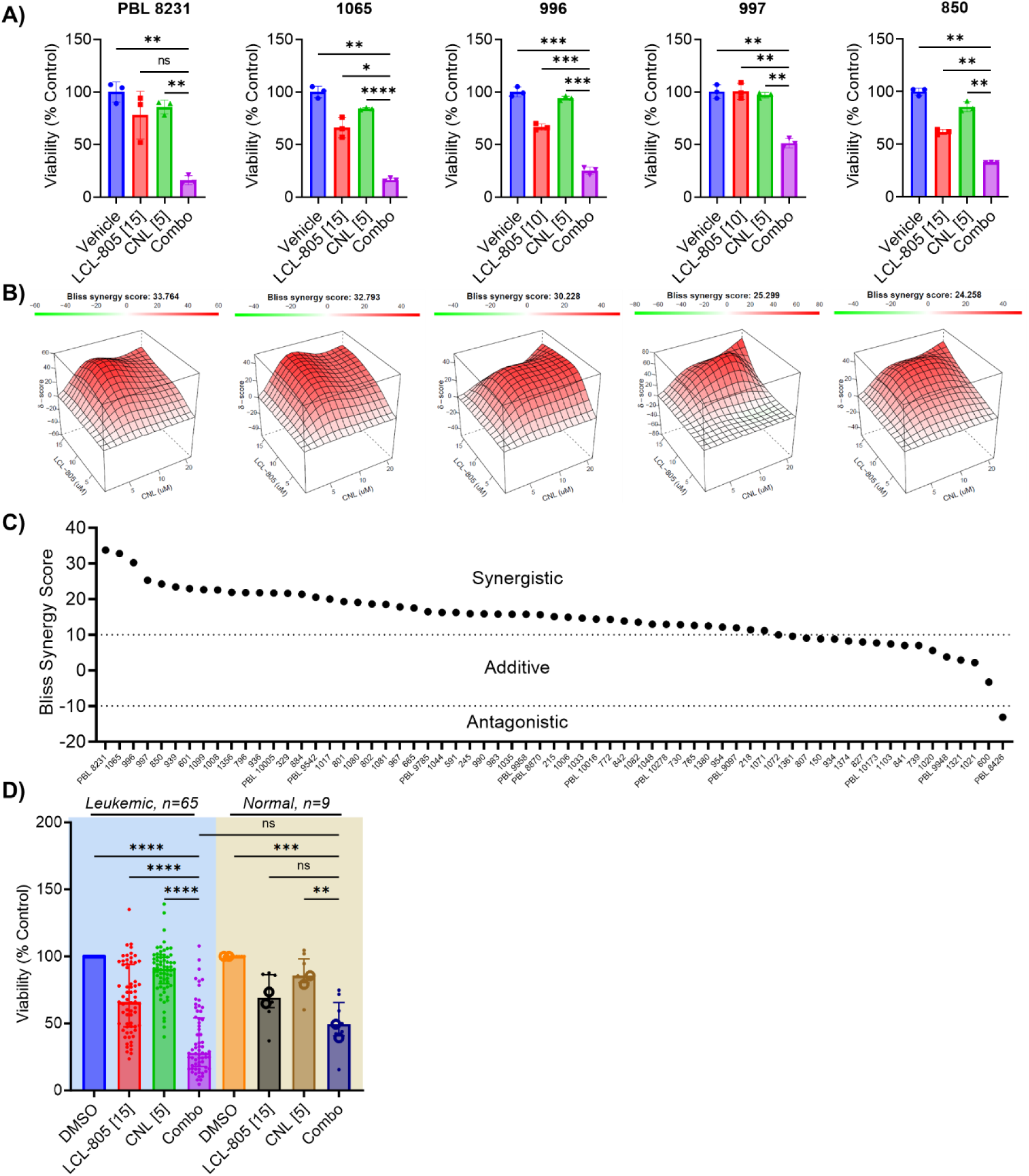
C6-ceramide nanoliposome (CNL) supplementation increased LCL-805 toxicity. Cell viability was measured in primary AML patient samples treated with multiple concentrations of LCL-805 (5, 10, or 15 µM) and CNL (5, 10, or 20 µM). (A) Cell viability and Bliss synergy scores (B) of the top five AML patient samples with the highest synergy scores treated with vehicle (DMSO), LCL-805, CNL or the combination. Drug concentration (in µM) is indicated in brackets within the x-axis labels. (C) Bliss synergy scores of primary AML patient samples treated with LCL-805 and CNL. (D) Cell viability of primary AML patient samples or normal controls (7 PBMC samples (•) and 2 CD34+ bone marrow samples (**o**)) treated with DMSO, LCL-805, CNL, or the combination. Drug concentration (in µM) is indicated in brackets within the x-axis labels. Bars represent the average of three technical replicates from a representative experiment. Error bars represent +/- standard deviation. Statistical analyses represent Welch ANOVA with Dunnett’s multiple comparisons test. * *p* < 0.05, ** *p* < 0.005, *** *p* < 0.001, **** *p* < 0.0001, ns = non-significant.

## 4. Discussion

In this work, we evaluated the anti-leukemic efficacy and cell death mechanisms of the dimethylglycine (DMG)-B-13 prodrug, LCL-805, in AML. B-13 is a ceramide analog and ceramidase inhibitor with demonstrated anti-tumor properties [19, 20]. The AC-targeting properties of B-13 were improved via the synthesis of lysosome-localizing DMG-B-13 prodrugs [21]. We showed that LCL-805 was an effective AC inhibitor and modulator of sphingolipid content with demonstrated anti-leukemic efficacy exacerbated by CNL supplementation *in vitro*. Our mechanistic studies showed that the cell death mechanisms of LCL-805 were not mediated through caspase-dependent apoptosis, lysosomal cathepsins, autophagy, or necroptosis. Instead, our findings revealed that LCL-805 toxicity was mediated by Akt antagonism and an iron-dependent form of cell death distinct from canonical ferroptosis.

LCL-805 is structurally related to another DMG-B13-prodrug, LCL-521. We compare our LCL-805 findings with published work evaluating the anticancer properties, sphingolipid modulating effects, and cell death mechanisms of LCL-521. LCL-521 has demonstrated anti-tumor efficacy and reduces sphingosine and S1P levels in cell line models of human breast adenocarcinoma [46]. A follow-up study evaluated the concentration-dependent effects of LCL-521 on AC protein levels and sphingolipid metabolites [47]. AC inactivation and decreased sphingosine content following LCL-521 treatment were reversible, which is consistent with our observed effects of LCL-805 on AC activation and sphingosine content in AML. Our observation of increased dihydroceramides following LCL-805 treatment is consistent with the finding that high concentrations of LCL-521 inhibit the enzyme responsible for converting dihydroceramides to ceramides, dihydroceramide desaturase-1 [47].

A separate study evaluated the mechanisms by which LCL-521 targeted myeloid-derived suppressor cells [32] and showed LCL-521-mediated cytotoxicity was blunted by inhibiting cathepsins B and D. In contrast, loss of cell viability was not rescued following cathepsin inhibition in our studies. We instead showed that Akt activation and iron chelation protected against LCL-805 cytotoxicity. LCL-805-mediated cell death was not rescued by the ferroptosis inhibitors liproxstatin-1 or ferrostatin-1 despite the protective effect of iron chelation. Iron chelation with DFO stabilized HIF-1α protein expression, which was further enhanced by LCL-805 treatment. HIF-1α stabilization promotes drug resistance in multiple leukemias, suggesting that HIF-1α stabilization protected against LCL-805 cytotoxicity [48]. These findings suggest that LCL-805 elicited a mode of iron-dependent cell death distinct from canonical ferroptosis. Differences in cell types evaluated between our study (human AML) versus the LCL-521 study (murine reticulum cell sarcoma) may contribute to the contrasting cell death mechanisms. Because lysosomes are a critical component of iron metabolism and AML cells are characterized by iron overload, the lysosome-targeting properties of LCL-805 may exacerbate an iron-dependent mode of cell death specific for AML [44, 49].

Multiple studies demonstrate that ceramide accumulation or the addition of exogenous ceramides antagonize Akt signaling [9]. In addition, others have linked AC to Akt-mediated cancer survival, drug resistance, and progression [41, 42]. We observed strong basal levels of Akt phosphorylation at Ser473 in MM-6 and OCI-AML2 cells. LCL-805 treatment dramatically reduced Akt phosphorylation. Supplementation with the Akt activator SC-79 protected against LCL-805 toxicity suggesting that Akt inactivation contributed to LCL-805 toxicity. In prostate cancer, AC overexpression promoted Akt activation involving sphingosine kinase 1 and S1P receptor 2 [42]. In our work, LCL-805-mediated S1P changes were modest, suggesting that the inhibitory effects of LCL-805 on Akt were likely mediated by elevated ceramide levels rather than loss of S1PR signaling.

The relationship between ceramide accumulation and cancer cell death provided rationale to combine exogenous ceramide with LCL-805 to enhance ceramide accumulation in the context of AC blockade. Previously, solubility and delivery issues due to the lipophilic nature of ceramides hindered exogenous ceramide supplementation as a viable treatment option. However, this major hurdle was overcome in part by the development of C6-ceramide-containing nanoliposomes (CNL) [10]. The LCL-805 and CNL combination resulted in potent synergistic lethality in primary AML patient samples warranting additional preclinical studies to define the efficacy and synergistic mechanisms of this combination. CNL completed a phase I trial in advanced solid cancers [50] and is currently entering a phase I/II trial as a single agent and in combination with vinblastine or with venetoclax and low dose Ara-C (LDAC) in relapsed/refractory AML. Sphingolipid metabolism and the regulation of mitochondrial apoptosis via Bcl-2 family proteins are intricately linked [51, 52]. The specific Bcl-2 inhibitor, venetoclax, is approved for use in AML in combination with LDAC or hypomethylation agents. However, high relapse rates from venetoclax-containing regimens provide the rationale and impetus to develop more effective venetoclax-based drug combinations. Ceramide-generating therapeutics in the form of exogenous ceramide [11] and sphingosine kinase inhibitors [53] improve the efficacy of venetoclax-containing AML regimens. These data highlight the utility of future studies combining AC inhibition with venetoclax.

The *in vivo* anti-leukemic efficacy of LCL-805 remains elusive as it exhibited toxicity and poor pharmacokinetic features. Thus, it failed to show efficacy in preclinical human cell line xenograft studies (data not shown). Future studies await improved nanoformulations or structural modifications of LCL-805 or its derivatives.

## 5. Conclusions

In this study, we demonstrated that LCL-805 inhibited AC activation to promote ceramide accumulation and cell death. LCL-805 was toxic towards AML cell lines and primary patient samples and synergized with C6-ceramide-containing nanoliposomes. Mechanistically, LCL-805 inhibited Akt signaling and induced iron-dependent cell death distinct from canonical ferroptosis. Because of its role in maintaining AML survival, AC remains a promising therapeutic drug target and further development of AC inhibitors is warranted.

## Acknowledgments

This work is dedicated in loving memory to Dr. Mark Kester, PhD. The authors thank those who provided cell lines for our studies: Dr. Francine Garrett-Bakelman, University of Virginia (F-36p and MOLM-16); Dr. Jacqueline Cloos and Carolien van Alphen, VU Medical Center Amsterdam (EOL-1, HEL, Kasumi-3, Kasumi-6, ME-1, ML2, MM-6 and NB4); Dr. Mark Levis, Johns Hopkins Medical Institutions (MOLM-13 and MOLM-14); Dr. Douglas Graham, Emory University (Nomo-1); Dr. Xiaorong Gu, Cleveland Clinic (OCI-AML2 and OCI-AML3); Dr. Harold L. Atkins, Ottawa Hospital Research Institute (OCI-AML4); Drs. Scott Kaufmann and Mithun Shah, Mayo Clinic (SET-2). We thank Galina Diakova and the UVA Biorepository and Tissue Research Facility (BTRF) (Research Resource Identifiers (RRID): SCR_022971) for providing primary AML patient samples and deidentified clinical data. The authors also acknowledge the contributions of the UVA ORIEN Team and the UVA BTRF in the consent of patients, specimen procurement, specimen processing, data abstraction, and providing access to molecular and clinical data (IRB-HSR #18445). We also thank Dr. James Norris, Medical University of South Carolina, for developing LCL-805.

## Author Contributions

- Conceptualization, JU, SFT, CEC, MCC, DJF, TPL
- Methodology, TEF; validation, JU, MT
- Formal analysis, JU, BJH
- Investigation, JU, JJPS, MT, UG, AS
- Resources, ZMS, HGW, DFC
- Writing—original draft preparation, JU, SFT, DJF
- Writing—review and editing, all co-authors
- Supervision, DJF, TPL
- Funding acquisition, MCC, TPL
- All authors have read and agreed to the published version of the manuscript.

## Institutional Review Board Statement

The study was conducted in accordance with the Declaration of Helsinki and approved by the Institutional Review Board of the University of Virginia (protocol code IRB-HSR #18445, 21625, 22123) and Penn State College of Medicine (Study ID: PRAMS029252EP, PRAMS040755EP).

## Informed Consent Statement

Informed consent was obtained from all subjects involved in the study.

---

**Figure S1.**
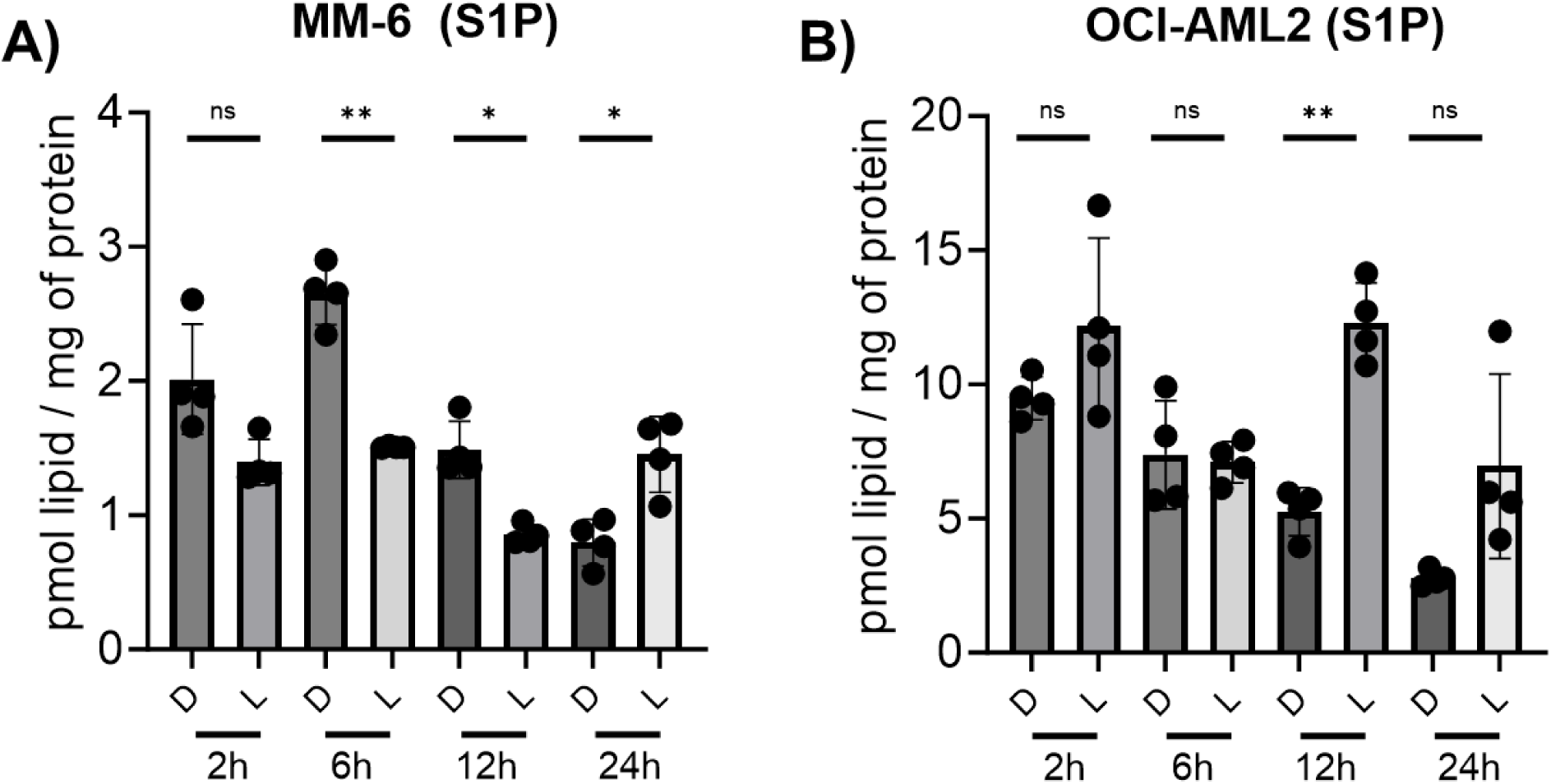
Effect of LCL-805 on sphingosine-1-phosphate levels. Sphingolipid profiling of MM-6 (A) and OCI-AML2 (B) cells treated with vehicle (DMSO) or LCL-805 (15 µM) for the indicated times. Levels of sphingosine-1-phosphate were evaluated by liquid chromatography mass spectrometry. Bars represent the average of four technical replicates from a representative experiment. Error bars represent +/- standard deviation. Statistical analyses represent Welch’s ANOVA with Dunnett’s T3 multiple comparisons test. * p < 0.05, ** p < 0.005, ns = non-significant. D = DMSO; L = LCL-805.

**Figure S2.**
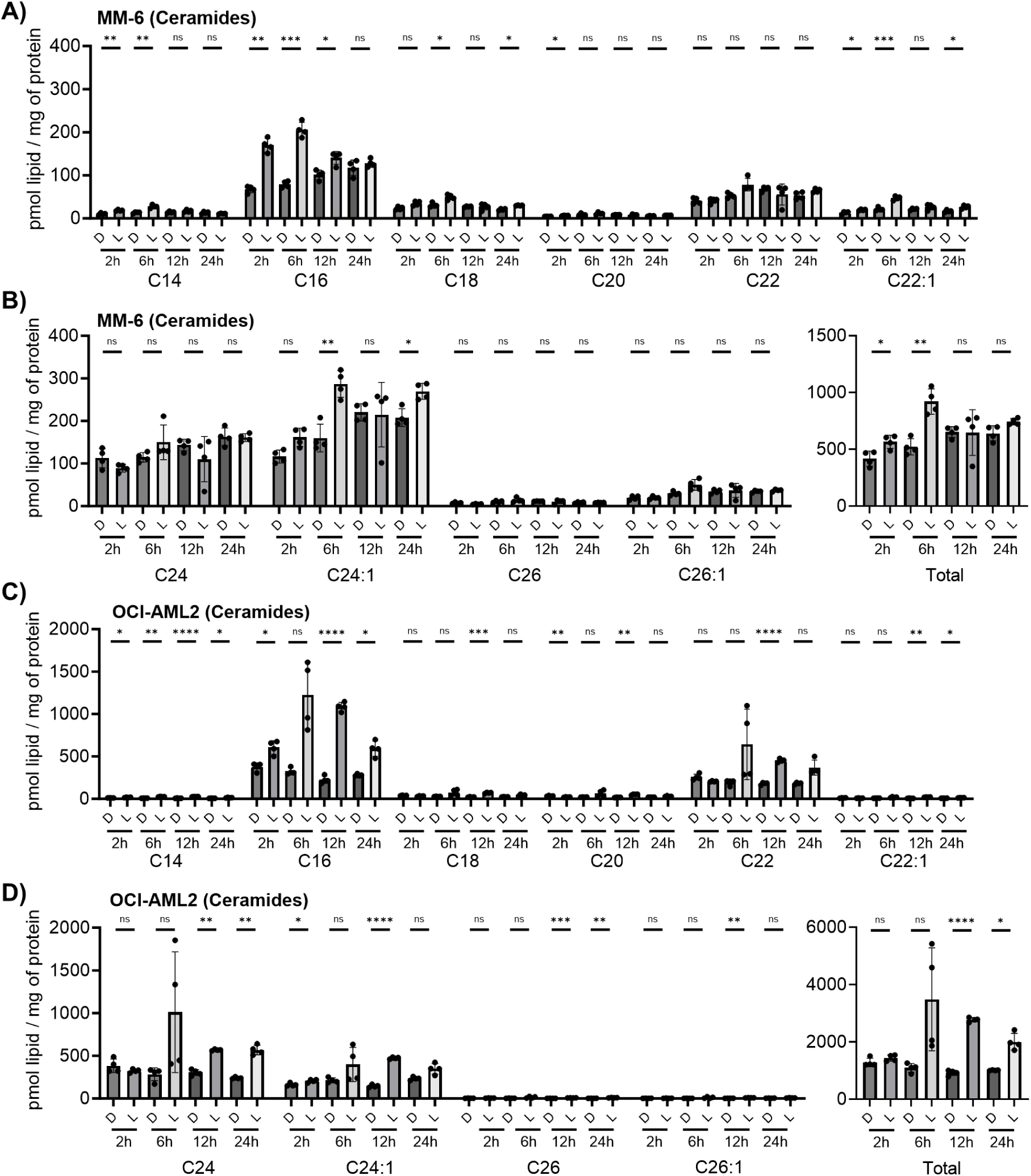
Effect of LCL-805 on ceramide levels. Sphingolipid profiling of MM-6 (A,B) and OCI-AML2 (C,D) cells treated with vehicle (DMSO) or LCL-805 (15 µM) for the indicated times. Levels of ceramide were evaluated by liquid chromatography mass spectrometry. Bars represent the average of four technical replicates from a representative experiment. Error bars represent +/- standard deviation. Statistical analyses represent Welch’s ANOVA with Dunnett’s T3 multiple comparisons test. * p < 0.05, ** p < 0.005, *** p < 0.001, **** p < 0.0001, ns = non-significant. D = DMSO; L = LCL-805. C## = fatty acid chain length. Total = sum of sphingolipid species.

**Figure S3.**
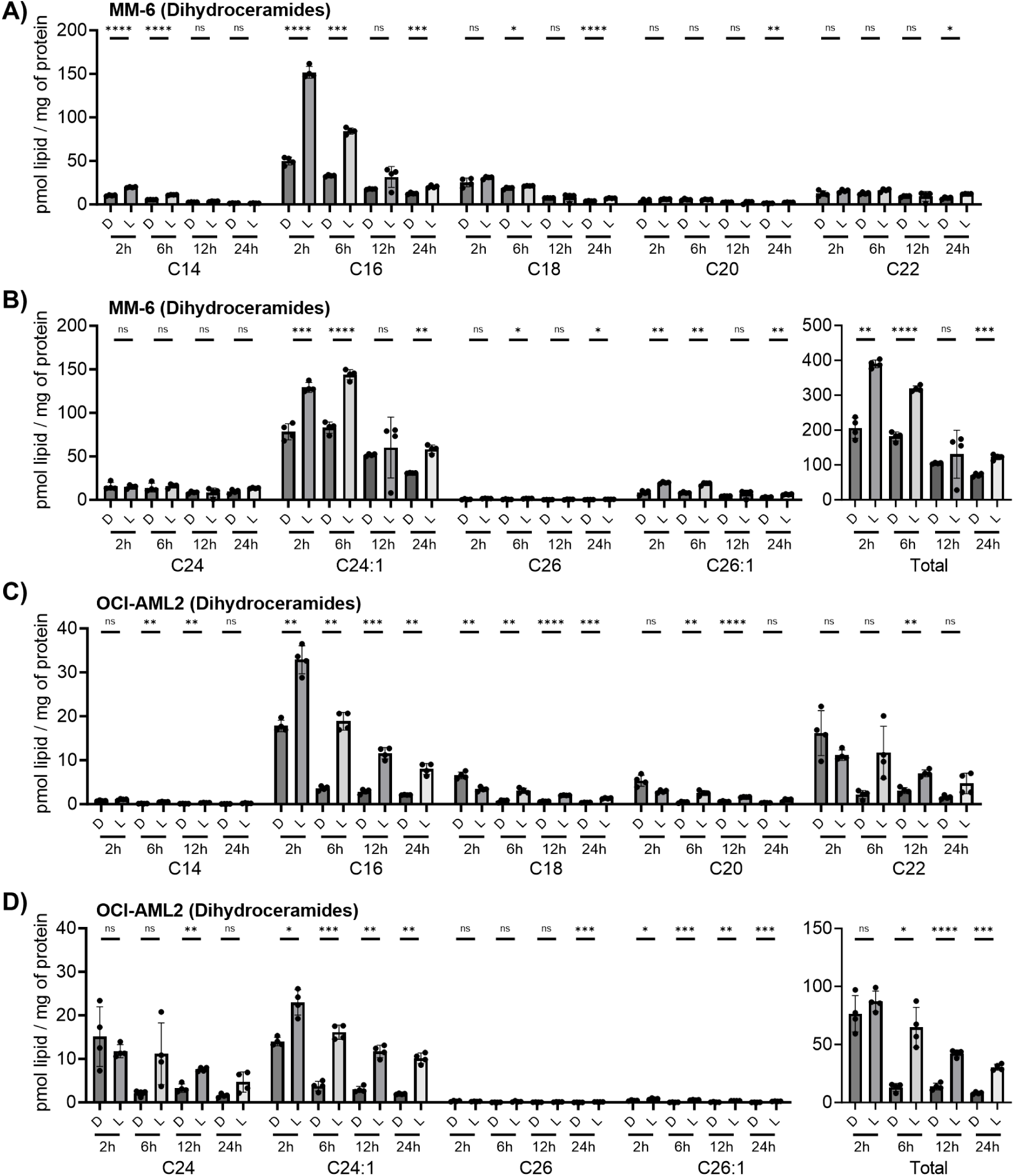
Effect of LCL-805 on dihydroceramide levels. Sphingolipid profiling of MM-6 (A,B) and OCI-AML2 (C,D) cells treated with vehicle (DMSO) or LCL-805 (15 µM) for the indicated times. Levels of dihydroceramide were evaluated by liquid chromatography mass spectrometry. Bars represent the average of four technical replicates from a representative experiment. Error bars represent +/- standard deviation. Statistical analyses represent Welch’s ANOVA with Dunnett’s T3 multiple comparisons test. * p < 0.05, ** p < 0.005, *** p < 0.001, **** p < 0.0001, ns = non-significant. D = DMSO; L = LCL-805. C## = fatty acid chain length. Total = sum of sphingolipid species.

**Figure S4.**
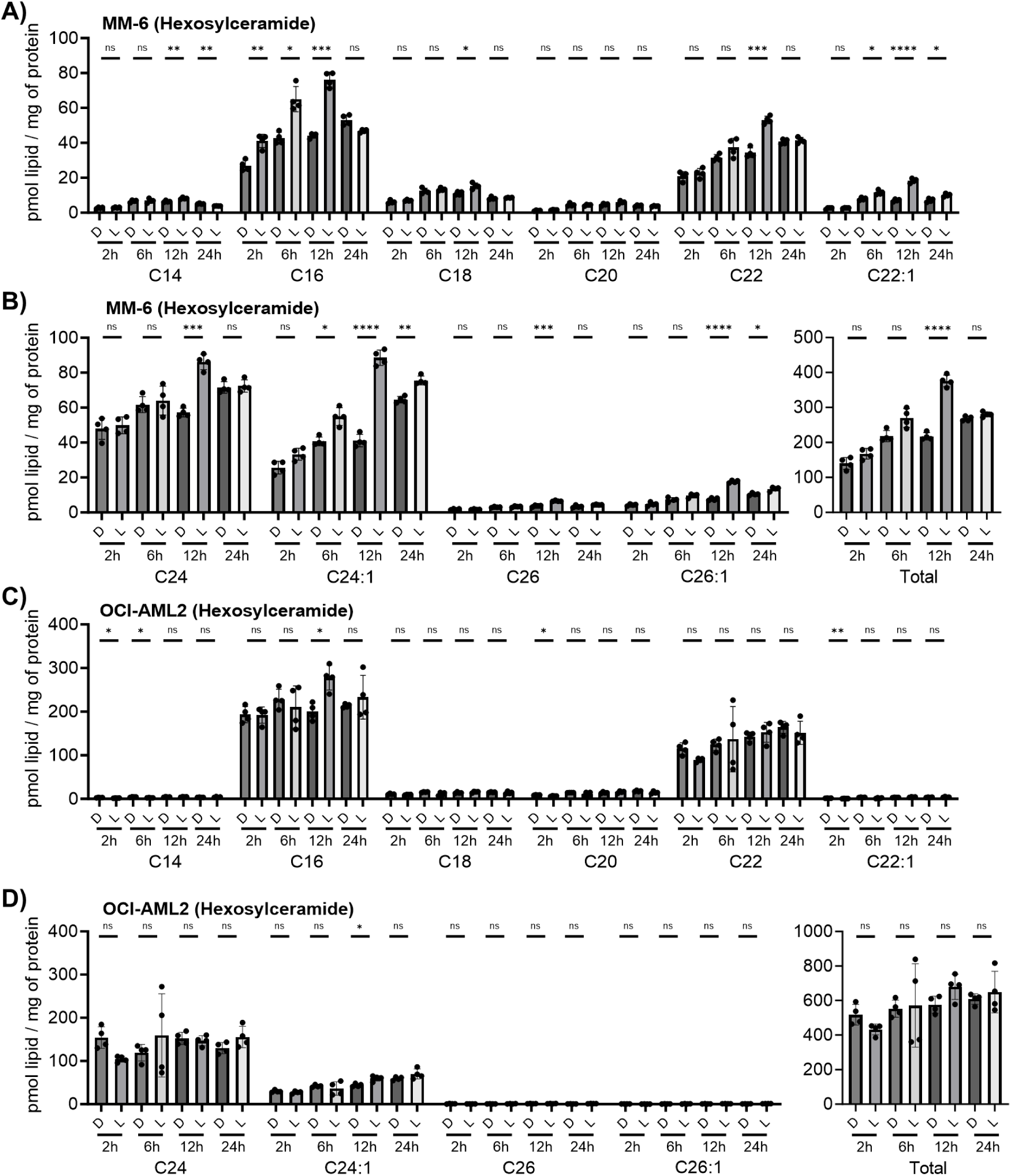
Effect of LCL-805 on hexosylceramide levels. Sphingolipid profiling of MM-6 (A,B) and OCI-AML2 (C,D) cells treated with vehicle (DMSO) or LCL-805 (15 µM) for the indicated times. Levels of hexosylceramide were evaluated by liquid chromatography mass spectrometry. Bars represent the average of four technical replicates from a representative experiment. Error bars represent +/- standard deviation. Statistical analyses represent Welch’s ANOVA with Dunnett’s T3 multiple comparisons test. * p < 0.05, ** p < 0.005, *** p < 0.001, **** p < 0.0001, ns = non-significant. D = DMSO; L = LCL-805. C## = fatty acid chain length. Total = sum of sphingolipid species.

**Figure S5.**
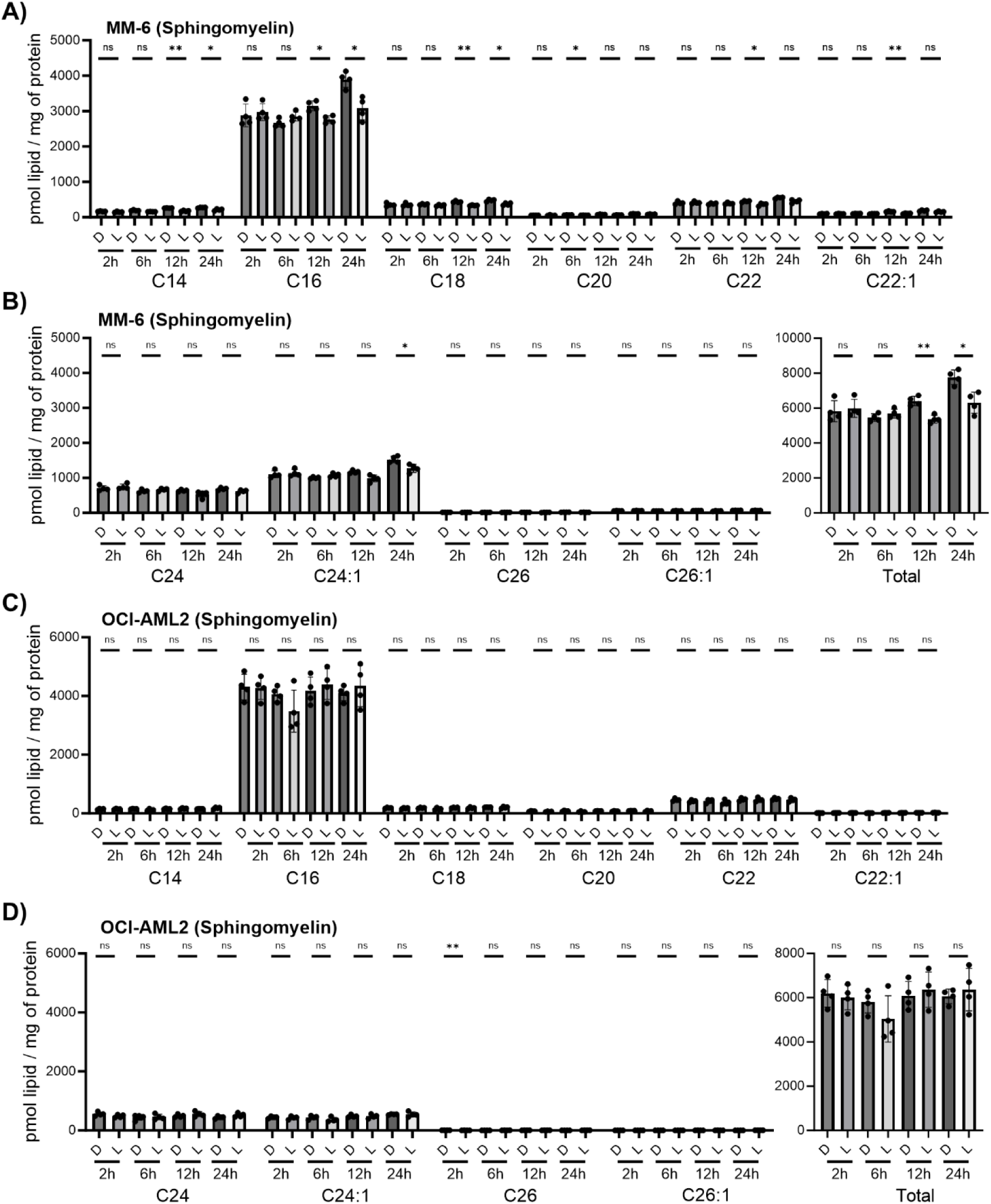
Effect of LCL-805 on sphingomyelin levels. Sphingolipid profiling of MM-6 (A,B) and OCI-AML2 (C,D) cells treated with vehicle (DMSO) or LCL-805 (15 µM) for the indicated times. Levels of sphingomyelin were evaluated by liquid chromatography mass spectrometry. Bars represent the average of four technical replicates from a representative experiment. Error bars represent +/- standard deviation. Statistical analyses represent Welch’s ANOVA with Dunnett’s T3 multiple comparisons test. * p < 0.05, ** p < 0.005, ns = non-significant. D = DMSO; L = LCL-805. C## = fatty acid chain length. Total = sum of sphingolipid species.

**Figure S6.**
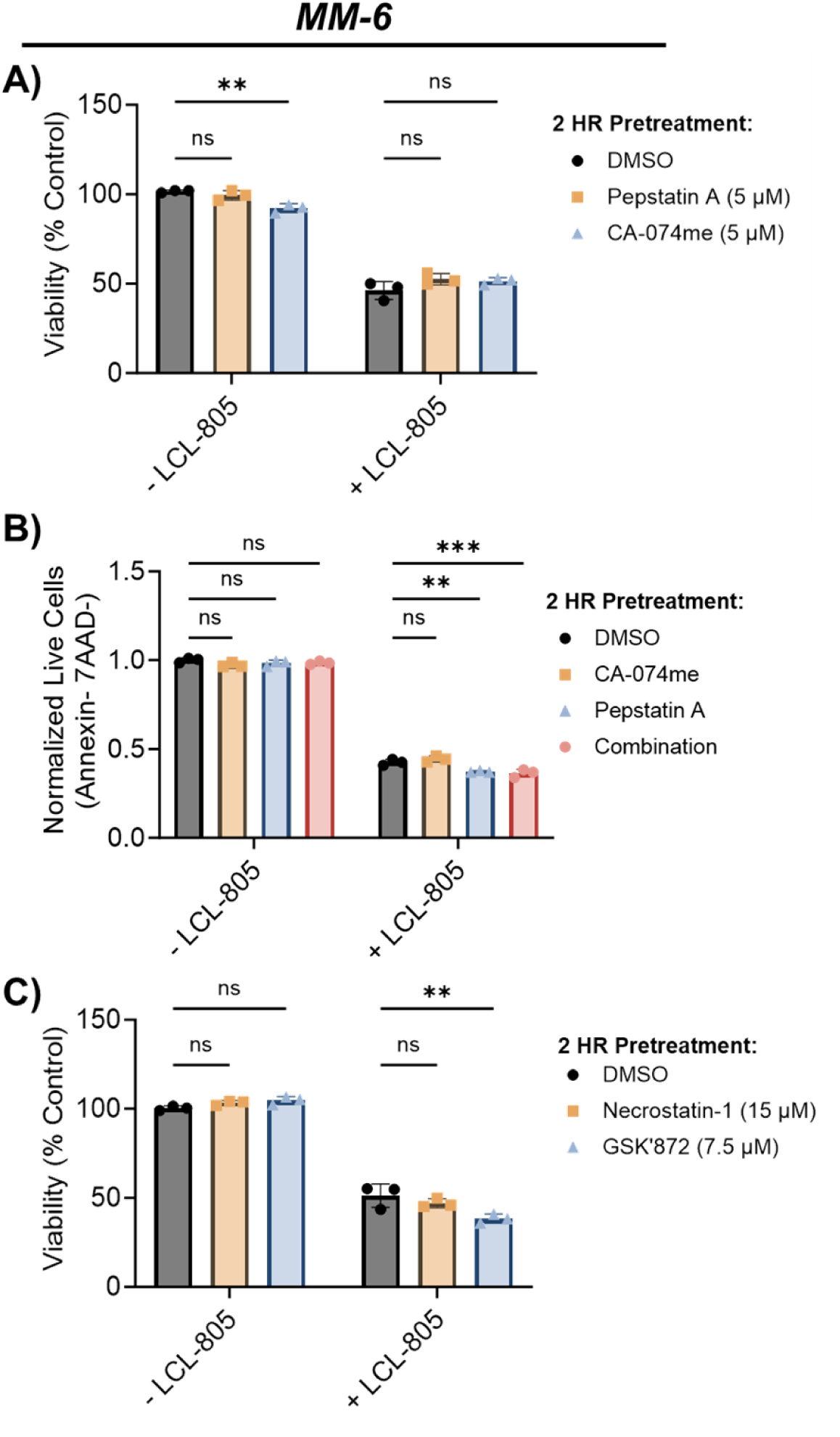
Inhibiting lysosomal cathepsins or necroptosis did not rescue LCL-805-mediated cytotoxicity. (A) Cell viability was determined by MTS assay of MM-6 cells pretreated with vehicle (DMSO), or lysosomal cathepsin inhibitors pepstatin A (5 µM), or CA-074me (5 µM) for 2 hours and subsequently treated with DMSO or LCL-805 (15 µM) for 24 hours. (B) Normalized live cell number by annexin and 7AAD flow cytometry of MM-6 cells pretreated with vehicle (DMSO), pepstatin A (5 µM), CA-074me (5 µM), or the combination for 2 hours and subsequently treated with DMSO or LCL-805 (15 µM) for 6 hours. (C) Cell viability was determined by MTS assay of MM-6 cells pretreated with vehicle (DMSO), or necroptosis inhibitors necrostatin-1 (15 µM), or GSK’872 (7.5 µM) for 2 hours and subsequently treated with DMSO or LCL-805 (15 µM) for 24 hours. Bars represent the average of three technical replicates from a representative experiment. Error bars represent +/- standard deviation. Statistical analyses represent two-way ANOVA with Tukey’s multiple comparisons test. ** *p* < 0.005, *** *p* < 0.001, ns = non-significant.

